# A universal regulatory mechanism for prevention of replication restart from RNA:DNA hybrids

**DOI:** 10.64898/2026.02.26.708313

**Authors:** Oyku Sensoy, Juan Carvajal-Garcia, Joshua R. Heyza, Paul A. Wiggins, Houra Merrikh

## Abstract

RNA:DNA hybrids form across all genomes. *In vitro* studies showed that these structures can be used to restart stalled replication forks, especially upon replication-transcription conflicts, however, this has not been tested *in vivo*. Here, we identify a mechanism that prevents replication restart from hybrids within cells. We identified the tRNA ligase, AsnRS, as a regulator of replication restart from hybrids at conflict regions in *Bacillus subtilis*. We determined that the DNA translocase Mfd and DNA polymerase PolA play key roles in this pathway. We unraveled a mechanism whereby Mfd removes RNA Polymerases, exposing 3’ ends of the RNAs within hybrids. These 3’ ends are then capped by AsnRS, preventing restart by PolA. Remarkably, the mammalian homologs of AsnRS, NARS1 and NARS2, fully complement the AsnRS phenotypes in bacteria. We propose that this is a universally conserved mechanism that prevents untimely replication initiation outside of origins from bacteria to humans.

## Introduction

RNA:DNA hybrids are found across genomes of all organisms. The nascent mRNA strand, which is synthesized by RNA Polymerase (RNAP), often hybridizes to the template DNA, displacing the non-template strand. This can occur at various stages of transcription, such as during RNAP backtracking, transcription termination, or simply during the elongation stage. Outside of Okazaki fragments, which are short-lived and form on the lagging strand of active replication forks, these structures lead to R-loop formation. *In vitro* studies have shown that mRNAs transcribed by RNAP can serve as primers to restart DNA replication upon stalling^1,2^. These studies were conducted using purified bacterial replication proteins. Given our collective knowledge of DNA replication *in vivo*, the results of these experiments led to a conundrum. Bacteria initiate replication from a single origin, once per cell cycle, and at the same time RNA:DNA hybrids form pervasively across the genome. Therefore, the results of these *in vitro* studies led us to the following question: why do we not observe replication initiation upon RNA:DNA hybrid formation elsewhere on the chromosome? This question extends to all species where replication origins are well-defined and RNA:DNA hybrids form across the chromosome.

Unlike in eukaryotes, replication restart is a well-characterized process in bacteria^3^. The replication machinery (replisome) encounters obstacles that stall its progression, including DNA binding proteins, DNA secondary structures, lesions on DNA, as well as and most commonly, RNAPs^4,5^. These stalling events leave various replication fork structures behind that are recognized by replication restart proteins to ensure the completion of genome duplication. Key players in replication restart include Pri proteins (e.g. PriA in *Bacillus subtilis* and PriA and PriC in *Escherichia coli*)^6^. PriA recognizes abandoned fork structures, and recombination intermediates (specifically D-Loops), after which it facilitates the recruitment of replisome proteins onto the DNA (i.e. DnaD, which loads the replicative helicase onto the DNA), allowing the replisome to reassemble and replication of the DNA to continue^7–11^. The classical replication restart mechanism, to our knowledge, does not depend on RNA:DNA hybrids.

Although replication and transcription are distinct biological processes, they both use the same DNA template, at the same time, leading to conflicts between the two machineries. Our group and others have found that there is an abundance of R-loops which by definition contain RNA:DNA hybrids at head-on conflict regions^12–15^. Interestingly however, we have not observed any replication restart at these regions and in fact, we found that the replication fork stalls indefinitely, at least under the conditions where we performed these experiments^12^. The *in vitro* studies mentioned above also examined the outcome of such conflicts using reconstituted replisome and transcription complexes. The results showed that the post-encounters between the replication and transcription machineries in either the head-on^2^ (genes encoded on the lagging strand) or co-directional^1^ (genes encoded on the leading strand) orientation, the nascent mRNA strand is used to restart replication. Therefore, the *in vitro* and *in vivo* studies in essence produced opposite results, further piquing our interest regarding RNA:DNA hybrid-driven replication restart.

The role of R-loops in replication has only been studied *in vivo* in the context of DnaA-independent replication initiation in cells with unresolved R-loops in the absence of RNase H, an endonuclease that degrades the RNA component of RNA:DNA R-loops and/or hybrids^16,17^. In these studies, rDNA-associated R-loops serve as replication initiation sites and allow DnaA and *oriC* independent replication initiation, which is known as constitutive stable DNA replication. It was proposed that in this special case, DNA polymerase I synthesizes DNA from the RNA primer within R-loops^18^. However, this suggestion in general or the role of DNA Polymerase I in this context has not been described and has simply remained as a hypothesis.

Our prior work demonstrated that unresolved RNA:DNA hybrids accumulate at sites of head-on replication-transcription conflicts^12^. While *in vitro* studies have shown that replication can restart using RNA:DNA hybrids, and *in vivo* studies in RNase H-deficient cells have demonstrated replication initiation outside the origin, our observations indicated a different outcome. In our system, unresolved RNA:DNA hybrids at sites of head-on replication-transcription conflicts lead to persistent replication stalling, inhibition of replication restart, and eventually, lethality^12^.

These findings led us to hypothesize that *in vivo*, there are active mechanisms that strictly prevent replication restart from RNA:DNA hybrids. To test our model, we first used several different engineered conflict systems inserted into the genome of *Bacillus subtilis*, where conflicts lead to excess RNA:DNA hybrids and determined the associated factors through pulldowns of the hybrids, followed by mass spectrometry (DRIP-MS). We identified AsnRS, an asparagine tRNA ligase, as a factor that physically associates with these hybrids at all sites of head-on replication-transcription conflicts that we engineered. Using genomic analyses, we found that AsnRS specifically prevents replication restart from RNA:DNA hybrids post-conflicts. Biochemical analyses revealed that AsnRS inhibits replication restart by binding to the 3’ ends of the RNA that are exposed once RNAP is removed by the DNA translocase Mfd and hybrids are exposed. We also found that AsnRS prevents DNA synthesis from mRNA primers by preventing DNA Polymerase I recruitment. Upon these findings, we examined mammalian cells, which have two homologs of AsnRS, NARS1 and NARS2. DepMap CRISPR gene-dependency analysis identified human cell lines with tolerance to NARS1 loss are enriched for DNA replication and replication stress response pathways. We introduced NARS1 and NARS2 into *B. subtilis* and observed full complementation of the phenotypic effects associated with the loss of AsnRS in the context of our study. This work shows that replication restart is regulated – a concept that has not been described. Furthermore, our work shows that like eukaryotes, prokaryotes also carry tRNA ligases that have moonlighting functions. Finally, our results strongly suggest that the replication restart mechanism described here is conserved across species. Overall, this study shows that cells have a safeguard against over-initiation outside of the origin, maintaining the correct DNA copy number across the genome while protecting genome stability.

## Results

### AsnRS associates with RNA:DNA hybrids at head-on conflict regions

We previously found that there is an abundance of RNA:DNA hybrids at head-on but not co-directional conflict regions^12^. As mentioned above, in contrast to the results of the *in vitro* studies^1,2^, we did not observe replication restart at head-on conflict regions. We hypothesized that cells may have mechanisms to regulate and/or inhibit replication restart form RNA:DNA hybrids. To investigate this model, we performed DRIPs after formaldehyde crosslinking, followed by quantitative mass spectrometry (Figure 1A). For this, we engineered three different *B. subtilis* cell lines that contained one of the following genes: *hisC*, *lacZ*, or the *luxA-E* operon (Figure 1B). In order to control for transcription, each gene was placed under an Isopropyl β-D-1-thiogalactopyranoside (IPTG) inducible promoter at the *amyE* locus. To control for gene orientation-dependent hybrid formation, we either inserted each gene on the leading strand (co-directional conflicts) or the lagging strand (head-on conflicts). To determine proteins associated with RNA:DNA hybrids, we used *rnhC* knockout cells, which lack the RNase HIII enzyme, allowing us to amplify the signal in these experiments by increasing the abundance of R-loops.

**Figure 1.**
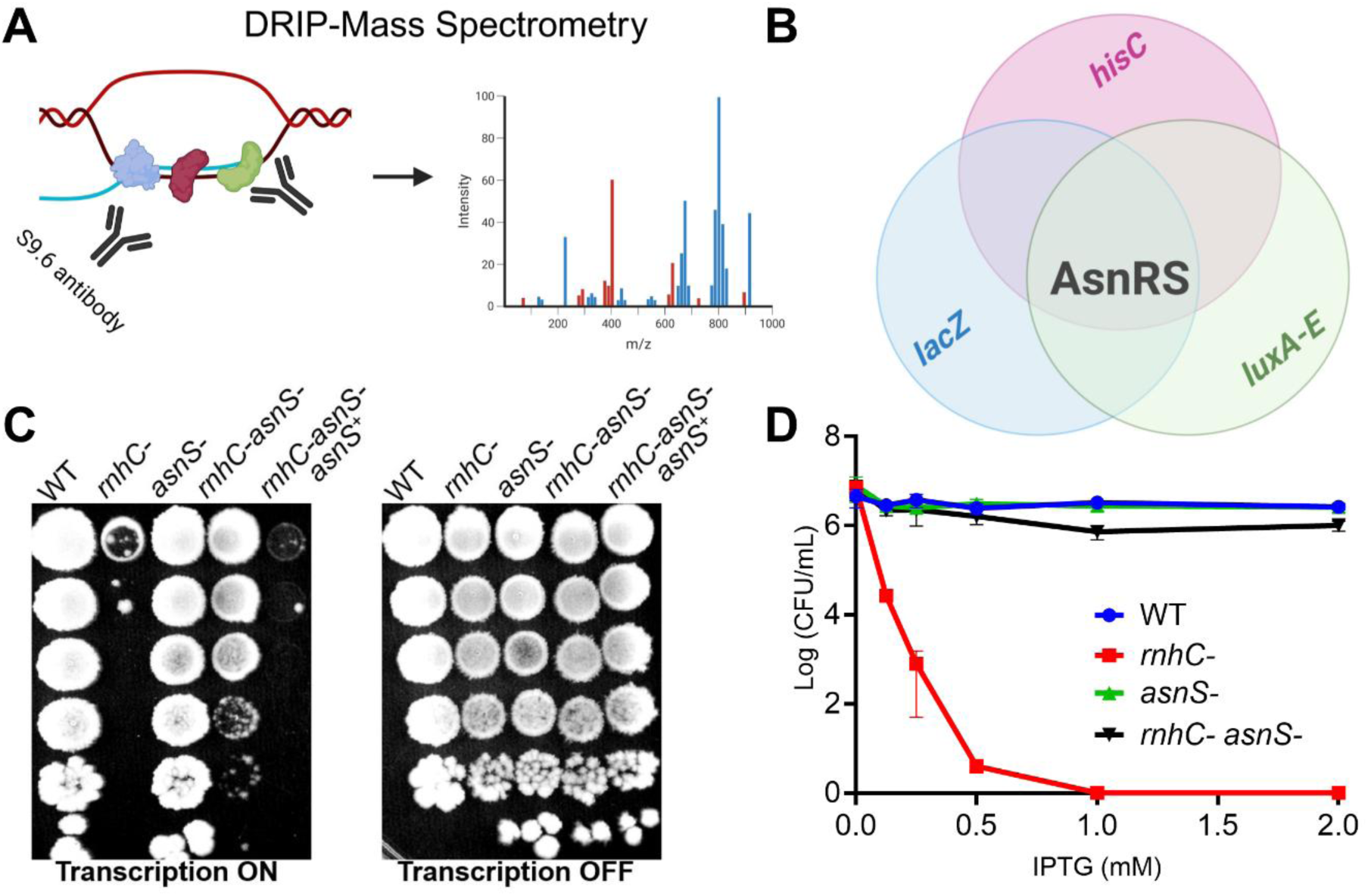
AsnRS, identified through DRIP-MS, inhibits cell growth upon head-on conflicts in cells without RNase HIII. A) Schematic of DNA:RNA immunoprecipitation followed by mass spectrometry (DRIP-MS) in *B. subtilis*. Following formaldehyde crosslinking, regions containing RNA:DNA hybrids and their associated proteins were immunoprecipitated using the S9.6 antibody. Proteins pulled down with RNA:DNA hybrids were then purified and identified by mass spectrometry. B) Venn diagram showing DRIP-MS identified AsnRS enriched in cells lacking RNase HIIII and experiencing engineered head-on conflicts created by inserting any of three reporter genes (*hisC, lacZ, luxA-E*) after filtration (Methods). C) Representative survival plates showing the growth of wild-type, *rnhC*-, *asnS*-, *rnhC*-*asnS*-, and *rnhC*-*asnS*-*asnS*+ (complemented by ectopic expression of *asnS* in cells lacking both RNase HIII and AsnRS) strains. All strains contain the reporter gene *hisC* at the engineered head-on conflict region. After spotting cells on LB plates supplemented with 1mM IPTG (left) and without IPTG (right), plates were incubated at 30°C overnight and removed from the incubator the next morning. Images were taken the following day. D) Survival assays were performed by inducing the reporter gene expression with IPTG in a dose-dependent manner (x-axis). Survival was measured by calculating CFU/ml for each strain spotted on plates (Figure 1C). Data are presented as the mean Log(CFU/ml) ± SEM from four independent experiments.

We identified proteins associated with both head-on and co-directional conflict regions in the presence and absence of RNase HIII. We examined each strain with and without IPTG to ensure that we are identifying proteins enriched upon transcription induction which is necessary for conflicts. We also performed several additional controls which can be found in Materials and Methods. We previously showed that with *lacZ* or *luxA-E* engineered conflicts there is severe replication fork stalling. However, we had not previously examined an engineered conflict system that utilized the *hisC* gene. To examine whether conflicts occur within the engineered head-on *hisC* gene, we performed DNA copy number analysis with and without RNase HIII in these strains. Relative copy number of each read was used as a proxy for replication progression. Similar to *lacZ* and *luxA-E*, the results showed uninterrupted replication at the head-on *hisC* locus in wild-type cells, while showing complete replication stalling in cells without RNase HIII when *hisC* was inserted in the head-on orientation (Supp. Figure 1A).

We also examined replication profiles of cells with the *hisC* gene on the leading strand, causing co-directional conflicts, and observed no replication stalling (Supp. Figure 1B). Furthermore, to rule out any global replication defect potentially occurring in the absence of RNase HIII, we examined replication progression in those cells without IPTG addition (Supp. Figure 1C). Both wild-type and RNaseH III-deficient cells exhibited uninterrupted replication profiles without the induction of head-on conflicts. These results further demonstrate that the replication stalling is due to head-on replication-transcription conflicts in the absence of RNase HIII.

We looked for factors that are associated with R-loops in all three strains, each with one of the head-on engineered conflicts, only when the genes were expressed via IPTG addition. We eliminated all the factors enriched in other strains tested (Materials and Methods). Applying all these criteria, we identified a protein that specifically associated with all three engineered conflict regions when the genes were oriented head-on to replication and RNA:DNA hybrids have accumulated at the head-on conflict regions: AsnRS (Figure 1B).

### AsnRS inhibits cell growth upon head-on conflicts in the absence of RNase HIII

AsnRS is a class II tRNA ligase that activates asparagine using ATP and attaches the activated amino acid to the 3’OH end of tRNA-Asn. Unlike most other tRNA ligases, AsnRS is not essential in all organisms due to a compensatory transamidation mechanism^19–23^. To investigate the role of AsnRS in head-on conflicts where R-loops and by proxy RNA:DNA hybrids are enriched, we assessed its impact on viability of cells in the presence and absence of RNase HIII while the cells were experiencing head-on conflicts. To do this, we performed survival assays on cells with engineered head-on conflicts by measuring colony forming units (CFUs) on Luria-Bertani media (LB) plates with varying concentrations of IPTG to induce expression of the reporter gene in a dose dependent manner, and quantified cell viability. In the absence of IPTG (“transcription off”), we did not observe any differences in viability across all four strains which included wild-type*, asnS-*, *rnhC-* and *rnhC- asnS-* double knockouts (Figure 1C and 1D). However, as we have demonstrated before, induction of transcription from the engineered conflicts obliterates survival in cells that are missing RNase HIII, only when the gene is expressed head-on but not co-directionally. Remarkably, the observed lethality in strains containing head-on conflicts, in the absence of RNase HIII, was largely reversed when we deleted the *asnS* gene (Figure 1C and 1D). Furthermore, we performed survival assays on cells with head-on engineered conflicts involving the other two genes from our initial DRIP-MS analysis (*lacZ* and *luxA-E*). We observed that the lethality was reversed in all cases in the absence of AsnRS ruling out any gene-specific role AsnRS might have (Supp. Fig. 2).

Given that the known function of AsnRS is in translation, we tested whether this effect is independent of translation. Removing the ribosome binding site (RBS) from *hisC,* preventing its translation at the conflict site, did not impact the results (Supp. Figure 3A,B). We observed the same outcome using the conflict system involving *lacZ* gene with and without RBS. Production of β-galactosidase (via x-gal) confirmed that translation occurred in the presence of RBS, yet AsnRS-deficient cells did not exhibit any global translation defects (Supp. Figure 3C). Like the previous constructs, RBS had no impact on the rescue of lethality in the absence of both RNase HIII and AsnRS. (Supp. Figure 3D). Together, these results demonstrate that the role of AsnRS in inhibiting growth during head-on conflicts with excessive R-loops is independent of translation.

### AsnRS does not exacerbate replication-transcription conflicts

It is possible that AsnRS exacerbates conflicts and therefore its absence rescues cell death. To test this hypothesis, we first measured RNAP association levels with the *hisC* conflict region using Chromatin Immunoprecipitation (ChIP), total mRNA levels using qPCR, and R-loop levels using DRIP-qPCR.

To assess RNAP enrichment, we performed ChIPs using an antibody specific to the β-subunit (RpoB) of RNAP. We did not observe any difference in RNAP enrichment at the head-on conflict regions in the various strains regardless of the presence or absence of RNase HIII and/or AsnRS (Figure 2A). Similarly, mRNA levels as measured by qPCR did not show any differences between *rnhC*- versus *rnhC*- *asnS*- double knockout cells. We previously showed that the levels of mRNA in *rnhC*- cells go down upon head-on conflicts, therefore these experiments produced results that are consistent with our previous findings (Figure 2B). Removing only AsnRS did not impact the levels of RNAP association or mRNA levels from the conflict regions compared to wild-type cells, suggesting that AsnRS does not directly affect transcription.

**Figure 2.**
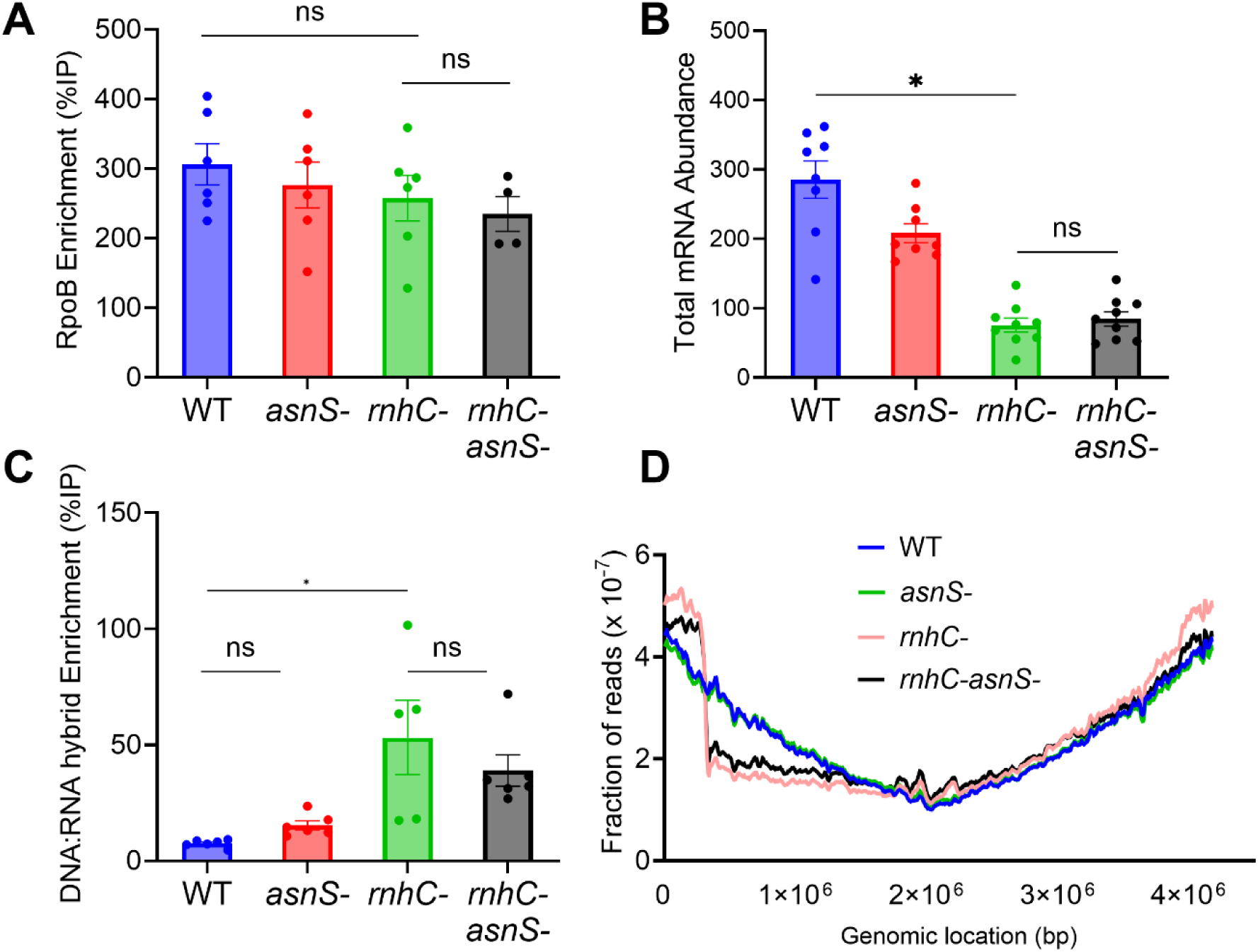
AsnRS does not exacerbate head-on replication-transcription conflicts. A,C) DRIP-qPCR and ChIP-qPCR analyses of wild-type, *rnhC*-, *asnS*-, *rnhC*-*asnS*- cells 2 hours after induction of engineered head-on conflicts by IPTG-induced expression of the reporter gene, *hisC*. RNA:DNA hybrids and RNA polymerase (RNAP) were immunoprecipitated using the S9.6 and RpoB antibodies, respectively. Enrichment of RNA:DNA hybrids and RNAP at the conflict region (*hisC*) was quantified by qPCR using the % input (%IP) method. Data are presented as mean ± SEM from three independent experiments (n = 9). Statistical analysis was performed using one-way ANOVA in GraphPad Prism; selected pairwise comparisons are indicated on the graph (ns, not significant; *p* < 0.05). B) Total mRNA transcript levels were measured in wild-type, *rnhC*-, *asnS*-, *rnhC*-*asnS*-cells 2 hours after induction of engineered head-on conflicts by IPTG-induced expression of the reporter gene, *hisC*. Following fixation with methanol (1:1), total RNA was extracted, reverse-transcribed into cDNA using random primers, and quantified by qPCR. Transcript levels corresponding to the engineered head-on conflict region (*hisC*) were quantified relative to a control region (*katA*). Data are presented as mean ± SEM from three independent experiments (n = 9). Statistical analysis was performed using one-way ANOVA in GraphPad; selected pairwise comparisons are indicated on the graph (ns: not significant, * p< 0.05). D) Marker frequency analysis was performed using whole-genome sequencing of wild-type, *rnhC*-, *asnS*-, *rnhC*-*asnS*- cells expressing the reporter gene (*hisC*) inserted into the genome in head-on orientation relative to replication for 1 hour. The frequency of each read was normalized to the total number of reads (y-axis) and mapped to the *B. subtilis* genome (x-axis).

To determine if R-loop levels were different between the four strains, we measured RNA:DNA hybrid levels at the conflict regions by performing DRIPs. Similar to the results of RNAP ChIPs and mRNA levels, we did not detect any significant difference in cells lacking only AsnRS compared to wild-type cells. Consistent with prior work, we found that RNA:DNA hybrid levels at the head-on conflict regions were significantly higher in RNase HIII-deficient cells compared to wild-type cells. Importantly, the increased RNA:DNA hybrids remained high in cells lacking both RNase HIII and AsnRS, strongly suggesting that conflicts occur in *asnS*- *rnhC*- double knockout cells (Figure 2C).

Our results from the RNAP and R-loop enrichment experiments, as well as mRNA levels, suggested that even though cells lacking both RNase HIII and AsnRS survive, they first experience the conflict. In other words, AsnRS does not seem to exacerbate conflicts or replication fork stalling. To directly test this, we performed genome marker frequency analysis after a short pulse (1 hour) with IPTG, which turns transcription on from the *hisC* gene, leading to conflicts in the head-on orientation. The results clearly showed that replication initially stalls at the head-on engineered conflict regions in the absence of *asnS* (Figure 2D).

### AsnRS prevents replication restart at head-on conflict regions

The results described above led us to hypothesize that upon conflicts, cells undergo a prolonged replication pause but ultimately restart replication at R-loop-rich, head-on conflict regions in the absence of AsnRS. We tested this model by performing marker frequency analysis again in the relevant strains, but this time looking at replication profiles over time. The results showed an increase in genome copy number within the cells lacking both RNase HIII and AsnRS, suggesting that replication restart is taking place in these cells, albeit slowly (Figure 3A and B). In fact, when we focus on the head-on conflict region, where replication stalls, we see that the DNA copy numbers double over time compared to the early time point in the same cells (Figure 3B).

**Figure 3.**
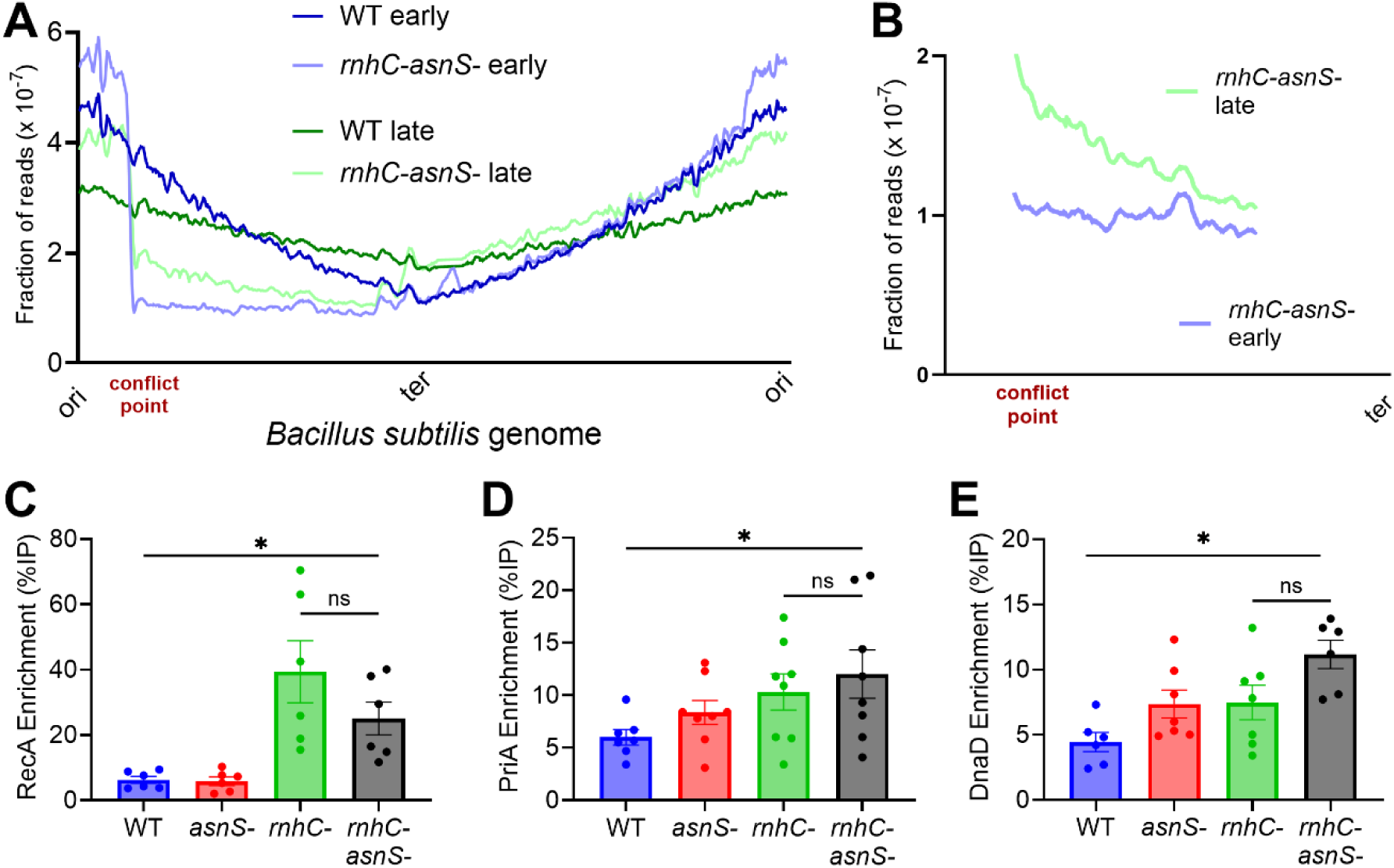
AsnRS prevents replication restart without impacting the enrichment of key replication restart proteins at head-on conflict regions in the absence of RNase HIII. A,B) Marker frequency analysis was conducted using whole-genome sequencing of wild-type, *rnhC*-, *asnS*-, *rnhC*-*asnS*- cells expressing the reporter gene (*hisC*) inserted into the genome in head-on orientation relative to replication (indicated as the “conflict point”) for 2 hours (early) and 7 hours (late). Read counts were normalized to the total number of mapped reads (y-axis) across the *B. subtilis* genome (x-axis). Panel A shows the genome-wide replication profile, and panel B shows a magnified view of the region surrounding the *hisC* insertion site. C-E) ChIP-qPCR analysis was performed to assess the associations of RecA, PriA, and DnaD, respectively, in wild-type, *rnhC*-, *asnS*-, *rnhC*-*asnS*- cells. Immunoprecipitation was performed using antibodies specific to each protein 2 hours after IPTG-induced expression of the reporter gene *hisC* at the engineered head-on conflict region. Enrichment of each protein at the conflict region (*hisC*) was quantified by qPCR using the % input (%IP) method. Data are presented as the mean ± SEM from at least two independent experiments (n=6). Statistical analysis was performed using one-way ANOVA in GraphPad; selected pairwise comparisons are indicated on the graph (ns: not significant, * p< 0.05).

However, given that when we turn on the head-on conflicts in a chronic manner on plates containing IPTG, cells lacking both RNase HIII and AsnRS grow similarly to wild-type in terms of CFUs, we expected to see a fully recovered DNA copy number in these double knockouts compared to the *rnhC-* cells. However, we did not observe a full recovery of DNA copy number as we anticipated during acute induction of conflicts over time. These results were perplexing; therefore, we turned to mathematical modelling and simulations.

We simulated bacterial marker frequency to predict differences in replication profiles with and without a replication pause. The results demonstrated that without a pause, there is a regular exponential decay toward the terminus, as seen in wild-type cells. Upon inducing conflicts, cells may display two different outcomes. They may exhibit a pause-like behavior which is temporary and shows a step-like profile. They may also show an arrest-like behavior that is a permanent pause and shows a flat profile (Supp. Figure 4). These predictions aligned with our observations indicating that while cells experience a replication arrest upon head-on conflicts in the absence of RNase HIII, their replication profile slightly differed from a flat form in the absence of AsnRS, suggesting a strong replication pause. Since we perform our experiments in asynchronous populations, we allowed enough time for each replication fork to encounter the conflict region. In this way we could observe the resulting behavior and compare it with our mathematical model. These predictions strongly correlated with our observations that cells lacking both RNase HIII and AsnRS exhibit a slow but step-like recovery in their replication profiles over time (Figure 3A, B).

Overall, both the *in vivo* data and the simulations indicated that cells lacking both RNase HIII and AsnRS can restart replication after pausing at head-on conflict regions. To determine how AsnRS prevents replication restart, we examined the recruitment of known replication restart proteins to the conflict region in the presence and absence of AsnRS. We measured the enrichment of key replication restart proteins PriA (which recognizes abandoned fork structures, recombination intermediates and replisome re-loading where replication forks have stalled and the replisome has dissociated from the replication fork)^6,11^, DnaD (the replicative helicase loader)^8^, and RecA (which generates D-loops by strand invasion, facilitating replication restart)^24^. We measured the enrichment of each protein at the head-on conflict regions by ChIP-qPCR. All three proteins showed increased enrichment at head-on conflict regions in the absence of both RNase HIII and AsnRS compared to wild-type cells. This increase did not change in cells without RNase HIII suggesting that AsnRS does not interfere with replication restart by preventing the recruitment of these critical replication restart proteins (Figure 3C-E).

### Replication restart from RNA:DNA hybrids requires PolA

Previous results showing the enrichment of some key replication restart proteins at the head-on conflict regions in the absence of RNase HIII, regardless of the presence or absence of AsnRS, suggested that perhaps AsnRS inhibits replication restart by influencing a downstream step. To test this model, we assessed whether PolA is required for restart from the hybrids *in vivo* in the absence of both RNase HIII and AsnRS. We chose PolA based on prior hypotheses^17,18^ that this DNA Polymerase may be responsible for any extension from RNA:DNA hybrids within R-loops (Figure 4A). To test PolA’s RNA-dependent DNA polymerase function, we performed *in vitro* primer extension assays. We used either DNA:DNA or RNA:DNA oligonucleotide substrates and allowed PolA to utilize the RNA primer to synthesize DNA upon the addition of dNTPs. Indeed, although significantly slower, PolA was able to synthesize DNA using both DNA and RNA primers hybridized to a longer DNA template *in vitro* (Figure 4B, Supp. Fig. 5). We then demonstrated that PolA is recruited to the head-on conflict regions by performing DRIP-MS in cells lacking both RNase HIII and AsnRS. The association of PolA with the head-on conflict regions depended on the absence of AsnRS, suggesting that AsnRS prevents PolA association with the RNA:DNA hybrids, inhibiting replication restart (Figure 4C).

**Figure 4.**
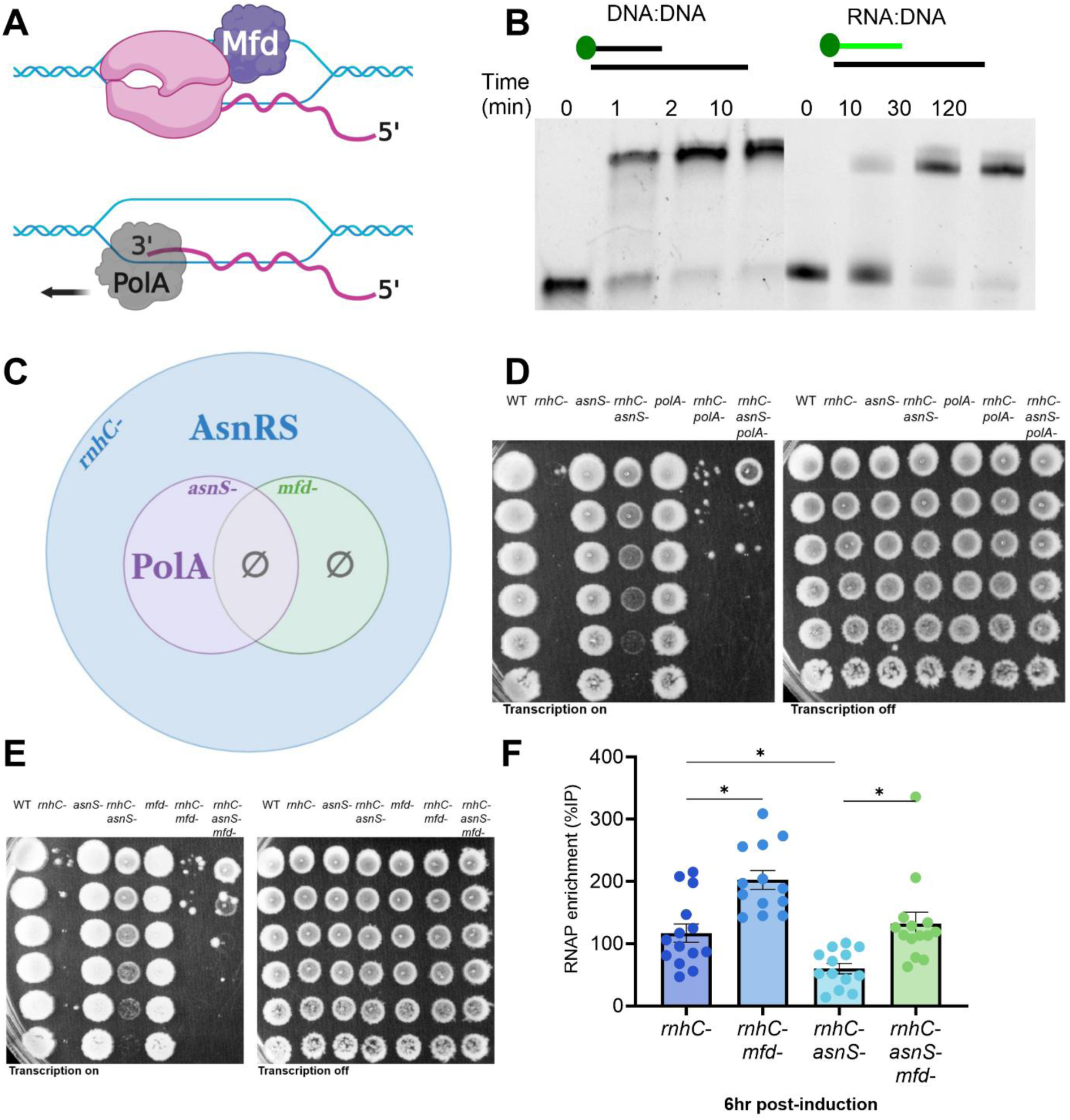
Replication restart from RNA:DNA hybrids at head-on conflict regions requires Mfd-mediated RNAP removal and PolA-dependent DNA synthesis. A) As proposed previously, PolA may restart replication from RNA:DNA hybrids by using the 3’OH end of the nascent RNA as a primer. For this extension to occur, the 3’OH end must be exposed; however, during transcription, it is localized within the active site of RNAP. Our model suggests that Mfd, a DNA translocase, displaces RNAP, freeing the 3’OH end of the nascent RNA to enable PolA function. B) Representative *in vitro* primer extension analysis quantifying DNA extension by PolA from DNA:DNA and RNA:DNA oligonucleotide templates using dNTPs. Samples were collected at the indicated time points, and reactions were stopped using 95% formamide–EDTA. Samples were analyzed on 10% TBE–urea gels. C) Venn diagram showing DRIP-MS, after filtration (Methods), identified AsnRS enriched in *rnhC*-, not in *rnhC-mfd-* cells while PolA enriched in *rnhC*-*asnS*- and not in *rnhC*- or *rnhC-asnS-mfd*- cells expressing the reporter gene (*hisC*) at the engineered head-on conflict regions. D,E) Representative survival assay plates showing the growth of corresponding strains, containing an IPTG-inducible gene (*hisC*) at the engineered conflict region. After spotting cells on LB plates supplemented with 1mM IPTG (left) and without IPTG (right), plates were incubated at 30°C overnight and removed from the incubator the next morning. Images were taken the following day. F) ChIP-qPCR analysis of *rnhC*-*asnS*-, *rnhC*-*asnS*-*mfd*- cells at a late time point (6 hours after IPTG-induced expression of the reporter gene *hisC* at the engineered head-on conflict region). RNAP was immunoprecipitated using a monoclonal antibody specific to RpoB. Enrichment of RNAP at the engineered head-on conflict region (*hisC*) was quantified by qPCR using the % input (%IP) method. Data are presented as the mean ± SEM from at least three independent experiments (n=9). Statistical analysis was performed using one-way ANOVA in GraphPad; selected pairwise comparisons are indicated on the graph (ns: not significant, * p< 0.05).

Unlike in *E. coli,* PolA is not essential in *B. subtilis* which allowed us to test its role phenotypically in this process. We performed survival assays (Figure 4D), and generated growth curves (Supp. Figure 6A) of cells with and without RNase HIII, AsnRS, and PolA. Removing PolA alone did not affect cell growth. Similarly, removing PolA in the absence of RNase HIII did not have any impact on lethality. However, removing PolA in the absence of both AsnRS and RNase HIII eliminated the rescue of the observed lethality, indicating that restart from the RNA:DNA hybrids depends on PolA.

### Replication restart from RNA:DNA hybrids requires Mfd

Replication restart eventually requires DNA extension from an RNA primer. In this case, the 3’OH end of the nascent mRNA would serve as a primer for DNA synthesis. However, given that the transcribed mRNA is likely to have remained inside RNAP upon stalling, such a process would require RNAP removal to expose the 3’ end for priming. The well-known DNA translocase and the anti-backtracking factor Mfd was an obvious candidate for facilitating this process^25^ (Figure 4A). Therefore, we hypothesized that RNAP is likely removed by Mfd to free the 3’ ends of the RNA, allowing PolA to initiate DNA synthesis in the restart process. Using similar approaches as described above, we tested whether Mfd is also required for replication restart in cells lacking both RNase HIII and AsnRS. Like PolA, we found that Mfd is also essential for replication restart under these conditions (Figure 4E, Supp. Fig. 6B). Importantly, when we looked at DRIP-MS of strains lacking Mfd, we did not find any enrichment for PolA at the conflict regions, largely solidifying that the removal of RNAP is required for the association of PolA with these regions (Figure 4C).

To further test this model (Figure 4A), we determined RNAP levels in both *rnhC*- and *rnhC-asnS-* cells, with and without Mfd. For this, we performed ChIPs of RpoB as previously described. We did not observe any significant difference in the presence and absence of Mfd when we measured the RNAP enrichment at an early time point (2 hours post induction) (Supp. Fig. 7). However, over time (6 hours post induction), RNAP levels increased in the absence of Mfd solidifying Mfd’s role in displacing stalled RNAPs (Figure 4F). Furthermore, RNAP levels stayed significantly higher in *rnhC*- cells compared to *rnhC-asnS-* cells. Together, these results show that both PolA and Mfd are required for replication restart, further supporting our model (Figure 4A).

### AsnRS prevents PolA from functioning through binding to RNA:DNA hybrids

Our results indicated that upon head-on conflicts with excess R-loops, AsnRS prevents replication restart, which requires Mfd and PolA. We investigated whether the catalytic activity of AsnRS is necessary for its role in interfering with replication restart. Based on the crystal structure solved in *Thermus thermophilus*, we identified the conserved residues within the ATP-binding site of AsnRS^26^. We mutated phenylalanine at position 219, which directly interacts with ATP, to alanine (Figure 5A)^27^. To determine whether this point mutation affected its catalytic activity, we performed a pyrophosphate assay measuring PPi production by AsnRS and AsnRS^F219A^. Indeed, the catalytic activity of the mutant we constructed was significantly impaired (Figure 5B).

**Figure 5.**
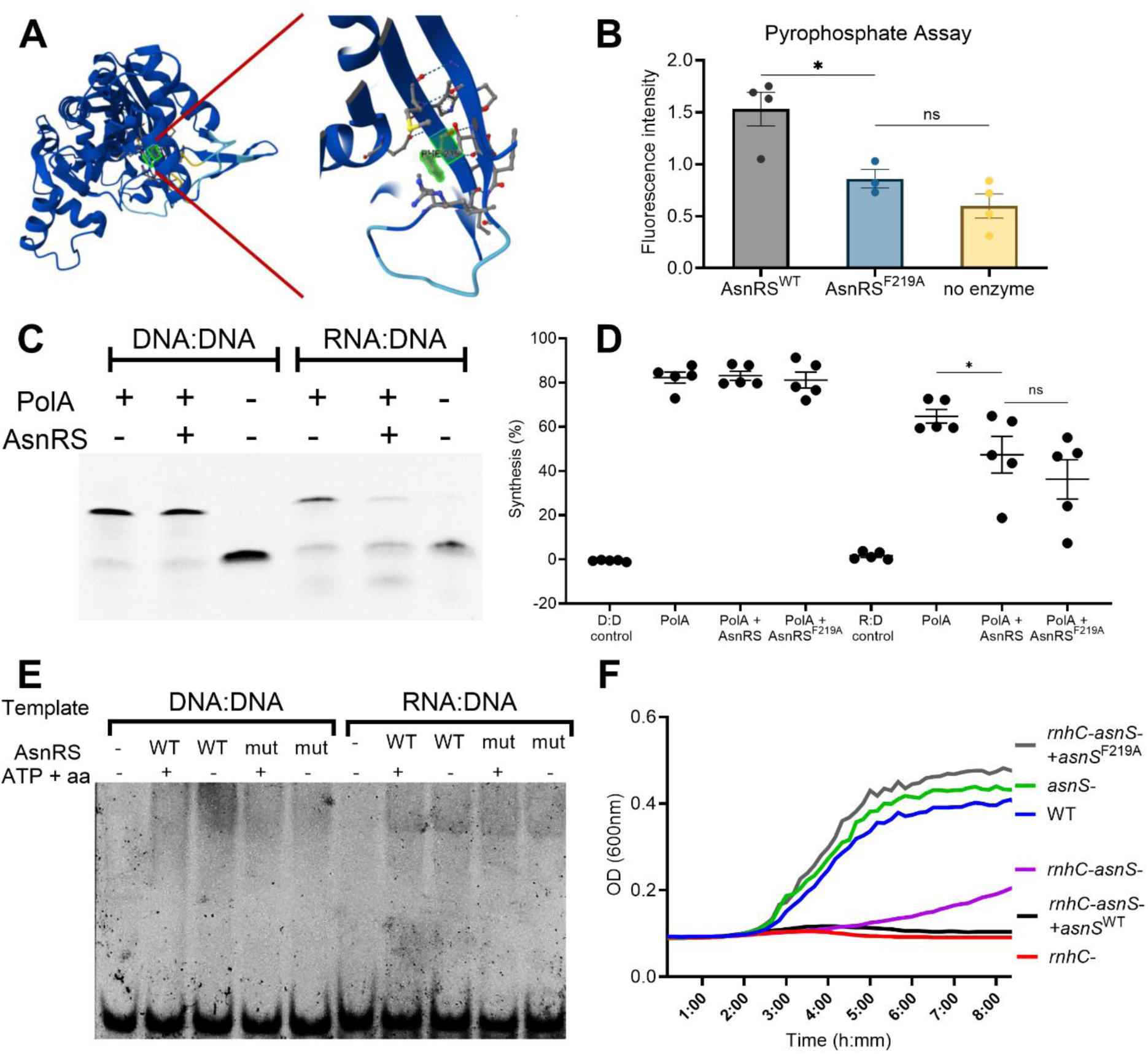
AsnRS binds RNA:DNA hybrids independently of its catalytic activity and inhibits PolA-mediated DNA synthesis. A) AlphaFold3-predicted the full-length structure of *B. subtilis* AsnRS (left)^27^. The zoom-in view of the ATP-binding site is also shown (right). Phenylalanine residue at position 219 (highlighted in green) was mutated to alanine (F219A) to impair AsnRS catalytic activity. B) Pyrophosphate assay was used to measure PPi production either by wild-type AsnRS or AsnRS^F219A^ in the presence of Asn, ATP, and yeast tRNA. Reactions were incubated at 37°C for 30 minutes and prepared for fluorescence measurement according to the manufacturer’s protocol. Fluorescence signal was measured by using BioTek NEO2 with Alpha Screening. Data are presented as the mean ± SEM from four independent experiments. Statistical analysis was performed using one-way ANOVA in GraphPad; selected pairwise comparisons are indicated on the graph (ns: not significant, * p< 0.05). C) Representative *in vitro* primer extension analysis quantifying DNA extension by PolA from DNA:DNA and RNA:DNA oligonucleotide templates in the presence and absence of AsnRS. Reactions were incubated with dNTPs for an hour at 37°C, and stopped using 95% formamide–EDTA. Samples were analyzed on 10% TBE–urea gels. D) *In vitro* primer extension assay was repeated using PolA in the presence of either AsnRS (wild-type) or AsnRS^F219A^. Band intensities were quantified using Image Lab software, and percent synthesis was calculated as the ratio of the extended product (upper band) to the total signal in each lane. Data are presented as the mean ± SEM from five independent experiments. Statistical analysis was performed using one-way ANOVA in GraphPad (ns: not significant, * p< 0.05). E) Representative Electromobility Shift Assay (EMSA) showing the binding of either AsnRS (wild-type) or AsnRS^F219A^ to DNA:DNA or RNA:DNA oligonucleotide templates. The templates were incubated in the presence of either protein with and without ATP (2mM) and asparagine (2mM) for 30min at 37°C. Samples were run on 5% polyacrylamide gels at 4°C. F) Growth curves of wild-type, *rnhC*-, *asnS*-, *rnhC*-*asnS*-, and cells complemented by ectopic expression of *asnS* and *asnS*^F219A^ in the absence of both RNase HIII and AsnRS (*rnhC*-*asnS*- +*asnS and rnhC*-*asnS*- +*asnS*^F219A^, respectively). All strains contain the IPTG-inducible reporter gene *hisC* at the engineered head-on conflict region and were grown in the presence of 1Mm IPTG. Data were plotted using the mean Optical Density (OD) values over 8 hours. Data are presented as the means of OD values from five independent experiments.

To determine whether AsnRS interferes with PolA-mediated DNA synthesis from RNA:DNA hybrids *in vitro*, we performed primer extension assays with and without AsnRS. While AsnRS had no impact on PolA-mediated extension from DNA primers, it inhibited the extension from the RNA primers (Figure 5C,D). The addition of asparagine did not impact the inhibition of PolA function by AsnRS (Supp. Figure 8). Furthermore, we observed that AsnRS^F219A^ similarly inhibited PolA function (Figure 5D, Supp. Figure 9). These results indicated that AsnRS inhibits PolA-mediated DNA synthesis from the hybrids and that this inhibition does not require AsnRS catalytic activity.

We examined whether AsnRS and/or AsnRS^F219A^ binds to either the DNA:DNA or RNA:DNA templates. To test this, we performed Electrophoretic Mobility Shift Assays (EMSA) and found that neither AsnRS nor AsnRS^F219A^ binds to the DNA:DNA template. However, both proteins bind to the RNA:DNA template regardless of the presence of ATP and asparagine (Figure 5E, Supp. Figure 10). Overall, these data indicate that AsnRS inhibits PolA-mediated DNA synthesis from RNA:DNA hybrids by binding to the RNA:DNA template, rather than through its catalytic activity, and that this template binding is sufficient to block PolA function.

We then examined whether the catalytically inactive mutant impacts the growth of cells. To test this, we complemented the double knockout cells, *rnhC-asnS*-, with either a wild-type copy of *asnRS or the asnRS^F219A^* mutant and observed that the wild-type AsnRS leads to loss of viability, whereas the AsnRS^F219A^ led to a full rescue of lethality (Figure 5F). These data suggest that the catalytic activity of AsnRS is harmful to cell growth. This result suggests that AsnRS has a secondary function outside of replication restart, a phenomenon that should be investigated but is outside the scope of our study.

### AsnRS’ role in regulating replication restart is not specific to engineered conflicts

Our engineered conflict system revealed that AsnRS is involved in regulation of replication restart during head-on conflicts when R-loops accumulate in the absence of RNase HIII. We next wanted to test whether AsnRS function in inhibiting replication restart is limited to engineered conflicts or dependent on the absence of RNase HIII. Many stress-response genes are oriented head-on to replication, and such stress-induced conflicts provide a biologically relevant setting to test whether AsnRS activity remains important even when RNase HIII is present during severe head-on conflicts.

We exposed wild-type and AsnRS-deficient cells to paraquat and lysozyme, two stressors that activate head-on genes (*katA* and *aphC*; *sigV* and the *dlt* operon, respectively)^12^. Remarkably, even with RNase HIII available to resolve RNA–DNA hybrids, loss of AsnRS improved cell survival under both stress conditions (Supp. Figure 11). These findings indicate that AsnRS-mediated inhibition of replication restart is not restricted to conflicts that are engineered or conditions lacking RNase HIII, but is important for naturally occurring head-on conflicts during exposure to stresses that induce head-on genes.

Another line of evidence supporting AsnRS role in replication restart comes from the genomic organization of *B. subtilis* and many other Gram-positive bacteria. The *asnS* gene, which encodes AsnRS, is within the same operon as *dnaD*^28^, encodes DnaD with a well-established role in replication initiation and restart. Given that genes localized within the same operon are often co-regulated and functionally related, this genomic arrangement suggests that AsnRS and DnaD may act in similar or cooperative pathways involved in maintaining replication integrity.

### Gene-dependency analyses suggest that the function of the human homolog of AsnRS, NARS1, is associated with DNA replication

To determine whether the replication-associated function of *B. subtilis* AsnRS is conserved in humans, we first examined gene dependency patterns for its human homolog, NARS1, across cancer cell lines using DepMap (Figure 6A). Across all profiled cell lines, NARS1 exhibited a strong negative gene effect score with a median of approximately −2 and is considered common essential in 1184/1186 cell lines, consistent with an essential role in cell viability (Figure 6A). However, two cell lines displayed modest tolerance to NARS1 loss, NCI-H1993 and HCC2998 with gene effect scores of −0.35 and −0.49, respectively. To identify genetic features associated with this resistance to NARS1 depletion, we performed a differential gene dependency analysis comparing NCI-H1993 to all other DepMap cell lines. This analysis revealed a set of 292 genes whose CRISPR dependency scores differed significantly in NCI-H1993 (q < 0.05) (Figure 6B). These results suggested that NCI-H1993 harbors genetic features that enable survival with reduced NARS1 function.

**Figure 6.**
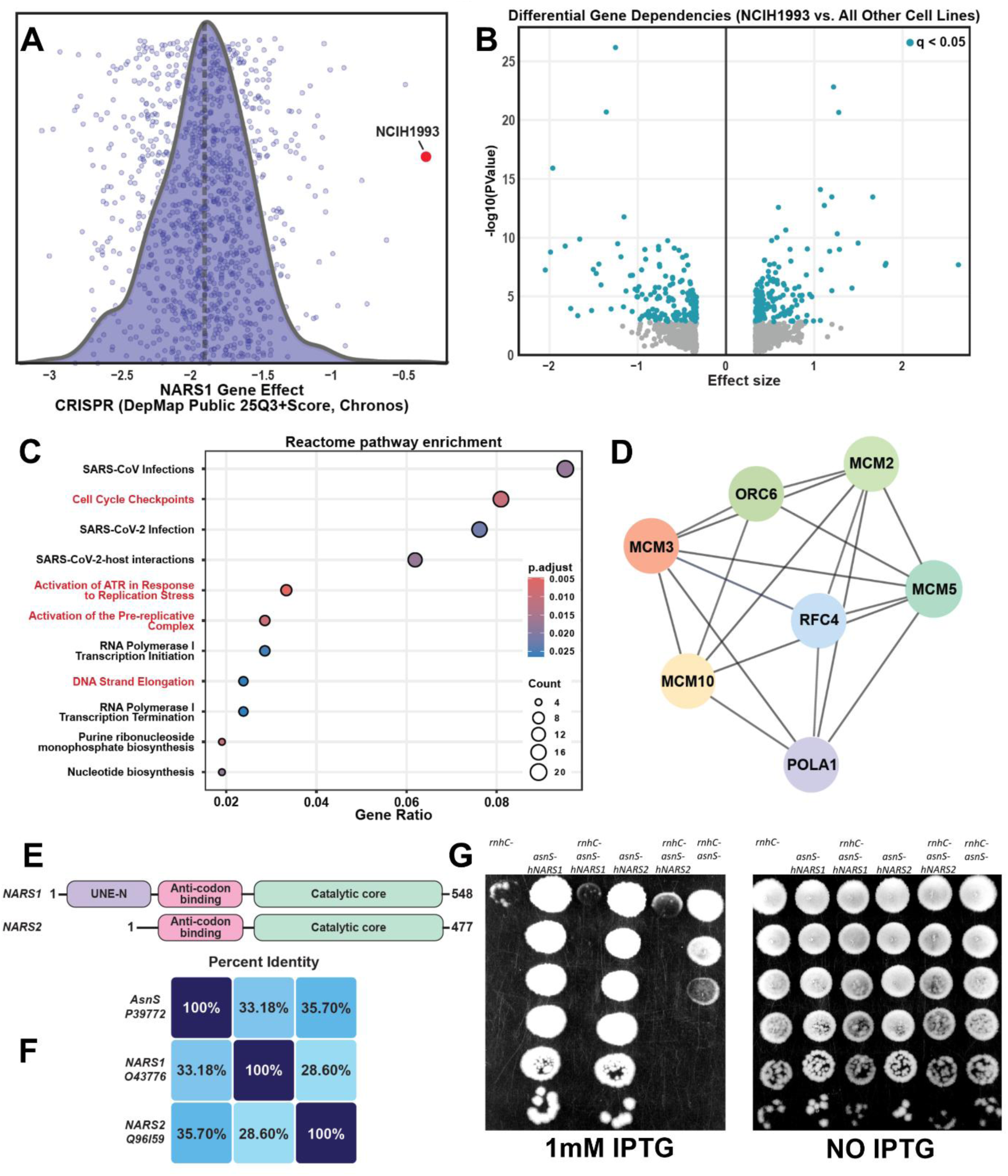
AsnRS/NARS defines a conserved replication-associated function across domains of life. A) Distribution of NARS1 gene effect scores from the DepMap Public 25Q3 CRISPR (Chronos) dataset across all profiled cancer cell lines. Individual points represent gene effect values for each cell line. B) Volcano plot depicting differential CRISPR gene dependency scores between the NCI-H1993 cell line and all other cell lines in the DepMap Public 25Q3 dataset. Each point corresponds to a gene. Genes with significantly different dependencies (q < 0.05) are highlighted in blue. C) Reactome pathway enrichment analysis of genes showing significantly different CRISPR dependencies in NCI-H1993 compared to all other DepMap cancer cell lines. D) STRING protein–protein interaction network of DNA replication–associated genes exhibiting significantly different CRISPR gene dependency scores in NCI-H1993 compared to other cancer cell lines. E) Schematic representation of the domain architecture of human NARS1 and NARS2. F) Pairwise percent amino acid sequence identity between *B. subtilis* AsnRS, human NARS1, and human NARS2. G) Representative survival assay plates showing growth of the indicated *B. subtilis* strains on LB plates supplemented with 1mM IPTG (left) or without IPTG (right). AsnRS deficiency in *B. subtilis* strains, *asnS*- and *rnhC-asnS*-, was complemented by ectopic expression of human *NARS1* or *NARS2* under an IPTG-inducible promoter. All strains also carry the IPTG-inducible reporter gene *hisC* at the engineered head-on conflict region. Cells were spotted onto LB plates with or without IPTG, incubated at 30 °C overnight, and imaged the following day.

**Figure 7.**
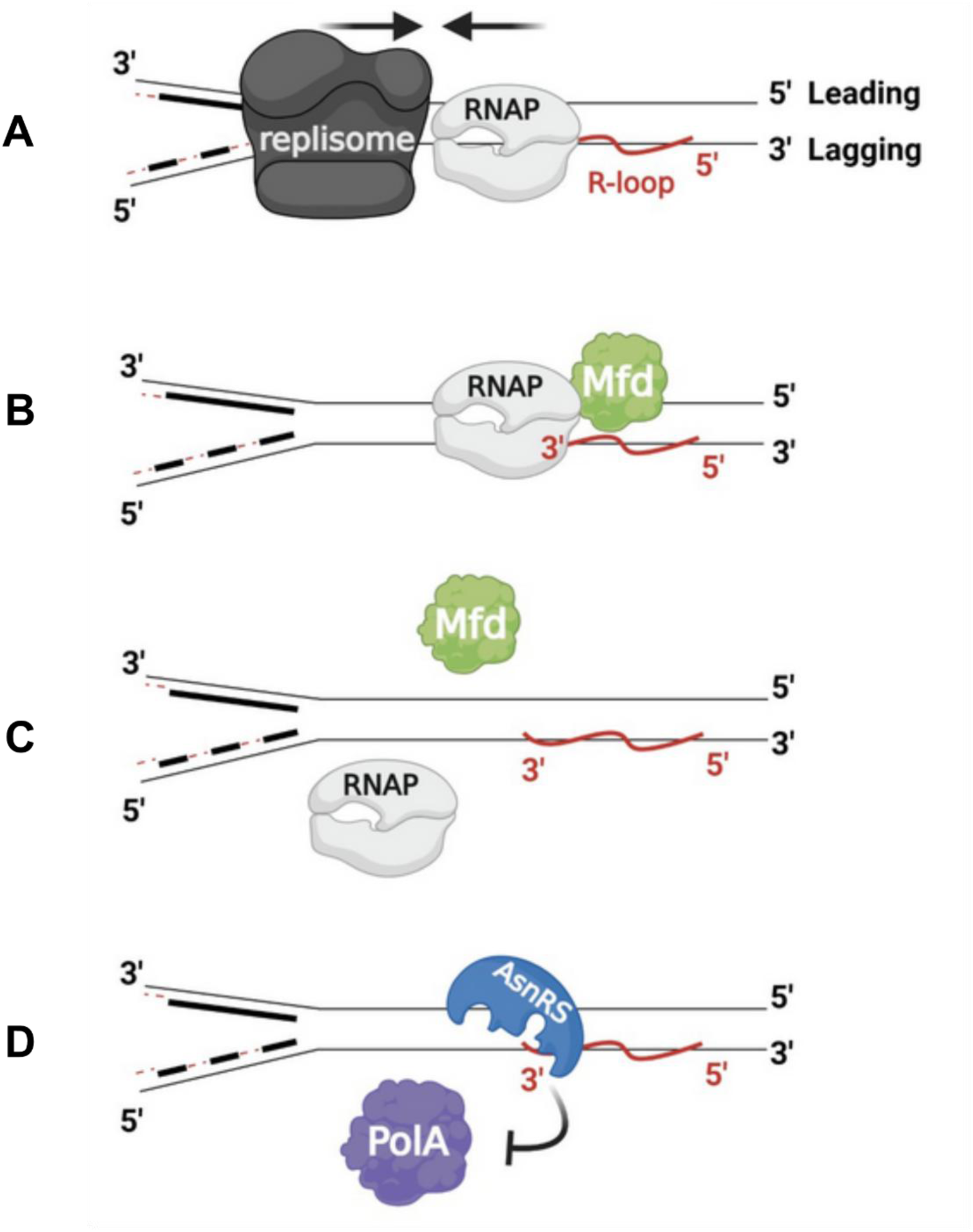
Working model of regulation of replication restart by AsnRS. A) When the replisome encounters RNA polymerase (RNAP) in a head-on orientation rather than co-directionally, cells accumulate R-loops in the absence of the R-loop resolution factor RNase HIII. These encounters stall both replication and transcription. B) Replisome stalling followed by disassembly necessitates a replication restart mechanism. During stalling, RNA:DNA hybrids that are formed co-transcriptionally. Transcripts produced at the conflict region can serve as primers for replication restart. However, the 3′-OH end of the nascent transcript, which is required to prime replication restart, remains captured by the stalled RNAP. Mfd, a bacterial DNA translocase and anti-backtracking factor, removes the stalled RNAP. C) Upon RNAP displacement by Mfd, the 3′-OH end of the RNA, is exposed. D) DNA polymerase I (PolA) can extend DNA using these RNA primers. E) However, AsnRS binds to the 3′-OH ends of RNA:DNA hybrids generated at head-on conflict regions, thereby blocking PolA-mediated DNA synthesis and inhibiting replication restart.

Next, we asked whether genes exhibiting differential dependency in NCI-H1993 were enriched for specific biological pathways. Reactome pathway enrichment analysis of the significant gene set revealed 11 significantly enriched pathways including DNA strand elongation, activation of the pre-replicative complex, activation of ATR in response to replication stress, and cell cycle checkpoints (Figure 6C). To further visualize functional relationships among these genes, we constructed a STRING protein–protein interaction network using replication-associated factors from the enriched gene set. This network revealed an interconnected module comprising core DNA replication components, including the catalytic subunit of Polα (POLA1), RFC4, ORC6, and multiple MCM complex subunits (MCM2, MCM3, MCM5, and MCM10) (Figure 6D). While RFC4, ORC6, and the MCM proteins had positive effect sizes in NCI-H1993 relative to other cell lines, POLA1 had an effect size of −0.88 indicating increased dependency of NCI-H1993 cells on POLA1. (Supp. Figure 12A). This is further supported by subsequent analysis in DepMap which showed that NCI-H1993 is among the most dependent cell lines on POLA1 expression (Supplemental Figure 12B). Notably, when both NCI-H1993 and HCC2998 cell lines were compared to all other cell lines, POLA1 remained a statistically significant hit (effect size: −0.56, q = 0.03986) indicating that this dependence is not unique to only NCI-H1993 cells.

### Human NARS1 and NARS2 functionally complement *B. subtilis* AsnRS deficiency

Eukaryotic genomes encode two AsnRS homologs: NARS1 and NARS2. NARS1 is suggested to localize to the cytoplasm, although whether it is also present in the nucleus is unknown. NARS2 is known to localize to mitochondria. Despite their distinct cellular compartments, both proteins retain the characteristic domain architecture of AsnRS enzymes, including an anti-codon binding domain and catalytic core (Figure 6E). Pairwise sequence comparisons revealed moderate amino acid sequence identity between *B. subtilis* AsnRS and human NARS1 (33.18%) or NARS2 (35.70%), as well as between the two human homologs themselves (28.60%) (Figure 6F), consistent with evolutionary conservation of core structural features and enzymatic functions.

To directly test whether human NARS proteins can functionally substitute for bacterial AsnRS, we codon-optimized NARS1 and NARS2 for expression in *B. subtilis* and introduced them into *B. subtilis* cells lacking AsnRS, with or without engineered conflicts and RNase HIII. We ectopically expressed either NARS1 or NARS2 in *rnhC-asnS-* cells to determine whether AsnRS function could be restored and thereby eliminate the rescue phenotype observed in the absence of AsnRS.

When we performed survival assays on these cells, we observed that AsnRS was complemented fully by both NARS1 and NARS2 (Figure 6G). Expression of either gene in cells lacking only AsnRS did not affect survival, ruling out potential toxic effects of human NARS1 or NARS2 expression in *B. subtilis* (Figure 6G).

## Discussion

In this work, we identify the first regulatory mechanism for replication restart^29,30^. Although restarting replication after the forks are stalled is necessary for cell survival (i.e. cells lacking PriA cannot survive)^11^, the regulation of this process seems to be critical for proper proliferation of cells when RNA:DNA hybrids are abundant. We identify a tRNA ligase, AsnRS, that seems to be a regulator of replication restart from RNA:DNA hybrids, which is the first example of bacterial tRNA ligases having a moonlighting function. Furthermore, although moonlighting functions of tRNA ligases have been observed in eukaryotic cells^31–36^, there has not been a connection described between these enzymes and DNA replication. The genomic organization of our model organism, *B. subtilis*, together with many other Gram-positive bacteria also highlighted potential function for AsnRS in replication restart since it is localized within the same operon as another crucial replication restart protein, DnaD. We also find that the mammalian homologs of AsnRS likely have a very similar function in DNA replication as to those described here. Overall, our studies provide insight into regulation of replication restart from the arguably most abundant restart substrate: RNA:DNA hybrids buried in R-loops.

In all organisms, regulation of replication initiation is tightly regulated. In bacteria, there are several regulatory functions for initiation from the origin, *oriC*, which include for example YabA in gram-positive^37,38^ organisms or SeqA in gram-negative^39,40^ species. Since there are so many opportunities in cells to restart replication, especially from RNA:DNA hybrids, it is quite reasonable that they would have tight regulatory mechanisms to prevent untimely DNA replication from regions that are not appropriate. Our work identifies the first of such regulatory mechanisms. AsnRS inhibits replication restart by preventing PolA reaching the 3’ end of the RNA primer within R-loops.

Our study was focused on regions where DNA replication and transcription complexes meet head-on, creating conflicts that involve the accumulation of R-loops (which include RNA:DNA hybrids). Given that AsnRS appears to specifically associate with these regions, we surmise that it is likely specific for prevention of restart during replication-transcription conflicts. However, this does not preclude AsnRS function at other regions of RNA:DNA hybrid enrichment which may have been under the limit of detection in our experiments.

We have published several studies where we propose that the head-on orientation for some genes may be beneficial for cells when it comes to adaptive evolution in bacteria^41–43^. This conclusion was based on several experimental studies as well as evolutionary analyses. When we measured mutation rates, we found that head-on conflicts lead to increased mutations^41^, and later, showed that this is due to R-loop formation^12^. Therefore, we propose that some tRNA ligases may be evolvability factors that influence mutagenesis.

Based on our results, we propose that RNAPs remain bound to R-loops that form co-transcriptionally^25,44,45^. As has been shown, these RNAPs are stalled, which triggers the association of the DNA translocase Mfd with the transcription complexes. The next stage in this process is that Mfd removes RNAPs as we have shown here supporting previous studies. This process exposes the 3’ end of the potential RNA primer.

Given that in several studies we have shown that Mfd, PolA, and R-loops are mutagenic^12,46,47^, this entire process may be of importance to evolution. This study associates tRNA ligases with a mutagenic and evolutionary pathway that we have described in detail in multiple studies. Perhaps AsnRS impacts mutation rates by prolonging RNA:DNA hybrids and/or regulating PolA function. This in itself is quite interesting, especially given the involvement of Mfd in this process. However, further investigation of this process is beyond the scope of this study.

We have shown that Mfd accelerates evolution and adaptation to antimicrobials^46,48^. Interestingly, cells lacking Mfd virtually have no phenotypic defects in the contexts where it has been studied: mainly during DNA damage and repair. Although Mfd influences mutagenesis, there have not been any studies showing that Mfd is essential for any cellular processes. The obliteration of survival in the triple mutant of *asnS- rnhC- mfd*-compared to *asnS- rnhC-* is the first major survival defect that has been associated with the absence of Mfd. This suggests that perhaps Mfd, which is thought of as a DNA repair factor (transcription-coupled repair), is essential when it comes to replication restart. This concept is novel and should be further investigated.

In our previous studies, we have shown that head-on conflicts are prevalent during stress conditions^12^. This is because most stress response genes are oriented head-on to replication and only get transcribed under conditions where cells are stressed, including in the presence of lysozyme, salt or reactive oxygen species that bacteria may encounter. In this study, we also showed that under such conditions, cells are better off (survive) in the absence of AsnRS. Therefore, AsnRS plays an important role in survival, in nature, where conditions can change rapidly and induce head-on conflicts.

Finally, the replication-associated function of AsnRS appears to be highly conserved. Using genome-scale CRISPR dependency data, we found that the human cytoplasmic AsnRS homolog, NARS1, is essential in nearly all cancer cell lines, yet a small subset of lines display partial tolerance to its loss. Genetic features associated with this tolerance in NCI-H1993 cells were enriched for DNA replication, replication stress signaling, and cell cycle checkpoint pathways. In particular, differential dependency analysis revealed altered dependencies in core replication initiation and elongation factors, including ORC, MCM complex subunits, RFC4, and Polα, suggesting that cells capable of surviving reduced NARS1 function may compensate for this loss through altered replication dynamics. Together, these observations support a conserved functional relationship between AsnRS/NARS proteins and replication restart pathways, despite the vast evolutionary distance between bacteria and humans.

Consistent with this interpretation, both human NARS1 and the mitochondrial homolog NARS2 were able to fully complement *B. subtilis* AsnRS deficiency. This functional complementation occurred despite only moderate sequence identity, indicating that the replication-regulatory activity of AsnRS is likely encoded within conserved structural or biochemical features rather than species-specific interactions. Consistent with our findings, a previous proteomic analysis of human RNA:DNA hybrid–interacting proteins identified NARS1 as an R-loop–associated factor^49^. Although that study did not explore a functional role for NARS1 in DNA replication or replication restart, its findings independently support a conserved association between AsnRS-family of enzymes and RNA:DNA hybrids. In addition, given the reliance of mitochondrial DNA replication on transcription-derived RNA primers, our findings raise the possibility that NARS2 may function as a regulator of mitochondrial replication restart or genome stability. Together with our functional and complementation data, these observations suggest that AsnRS-mediated regulation of RNA:DNA hybrid associated processes is likely evolutionarily conserved, from bacteria to humans.

## Supplementary figure legends

**Supp. Figure 1.**
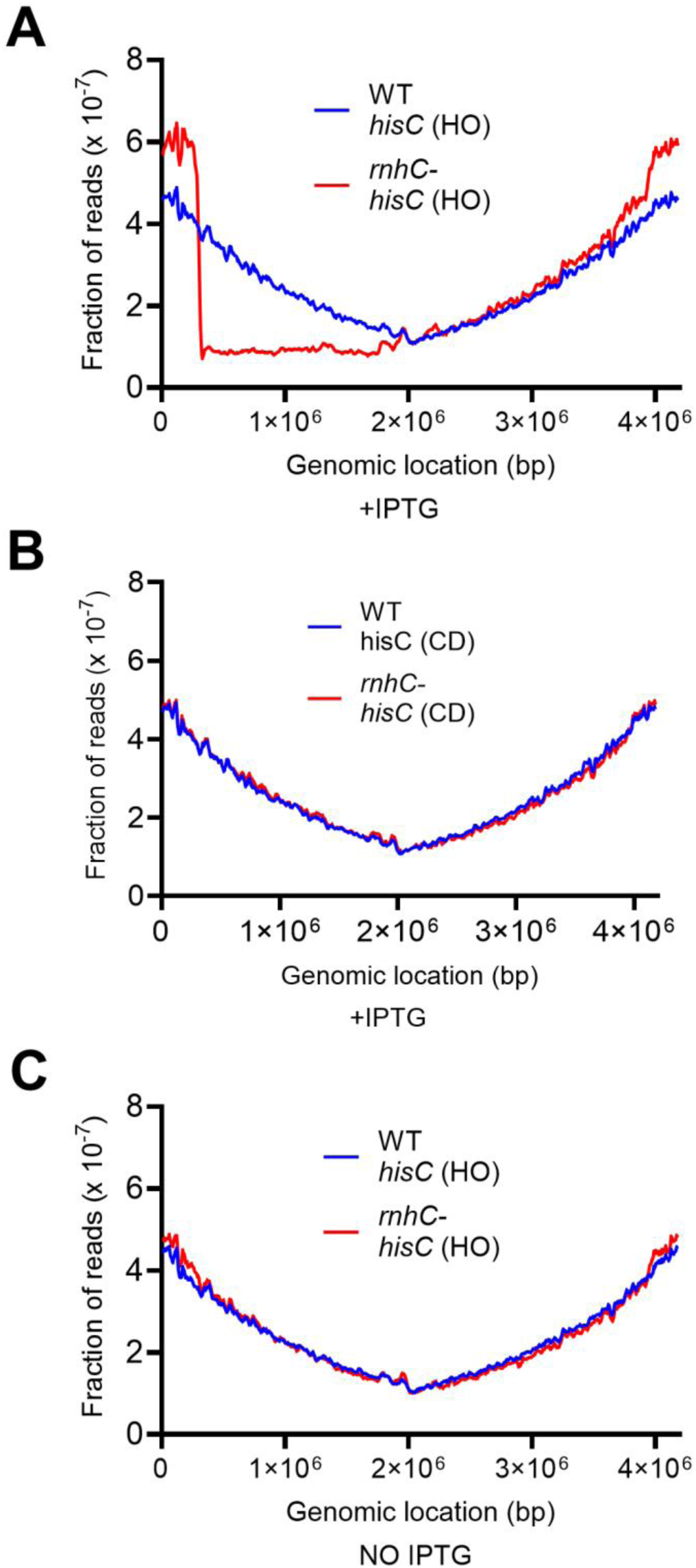
Replication stalls where head-on replication-transcription conflicts occur in the absence of RNase HIII. Marker frequency analysis was performed using whole-genome sequencing of wild-type and *rnhC*- cells expressing the reporter gene (*hisC*) inserted into the genome in head-on (A,B) or co-directional (C) orientation relative to replication. Cells were grown in the presence of IPTG, experiencing engineered conflicts in either head-on (A) or co-directionally (C), or without IPTG (B) for 2 hours followed by DNA extraction and sequencing. The frequency of each read was normalized to the total number of reads (y-axis) and mapped to the *B. subtilis* genome (x-axis).

**Supp. Figure 2.**
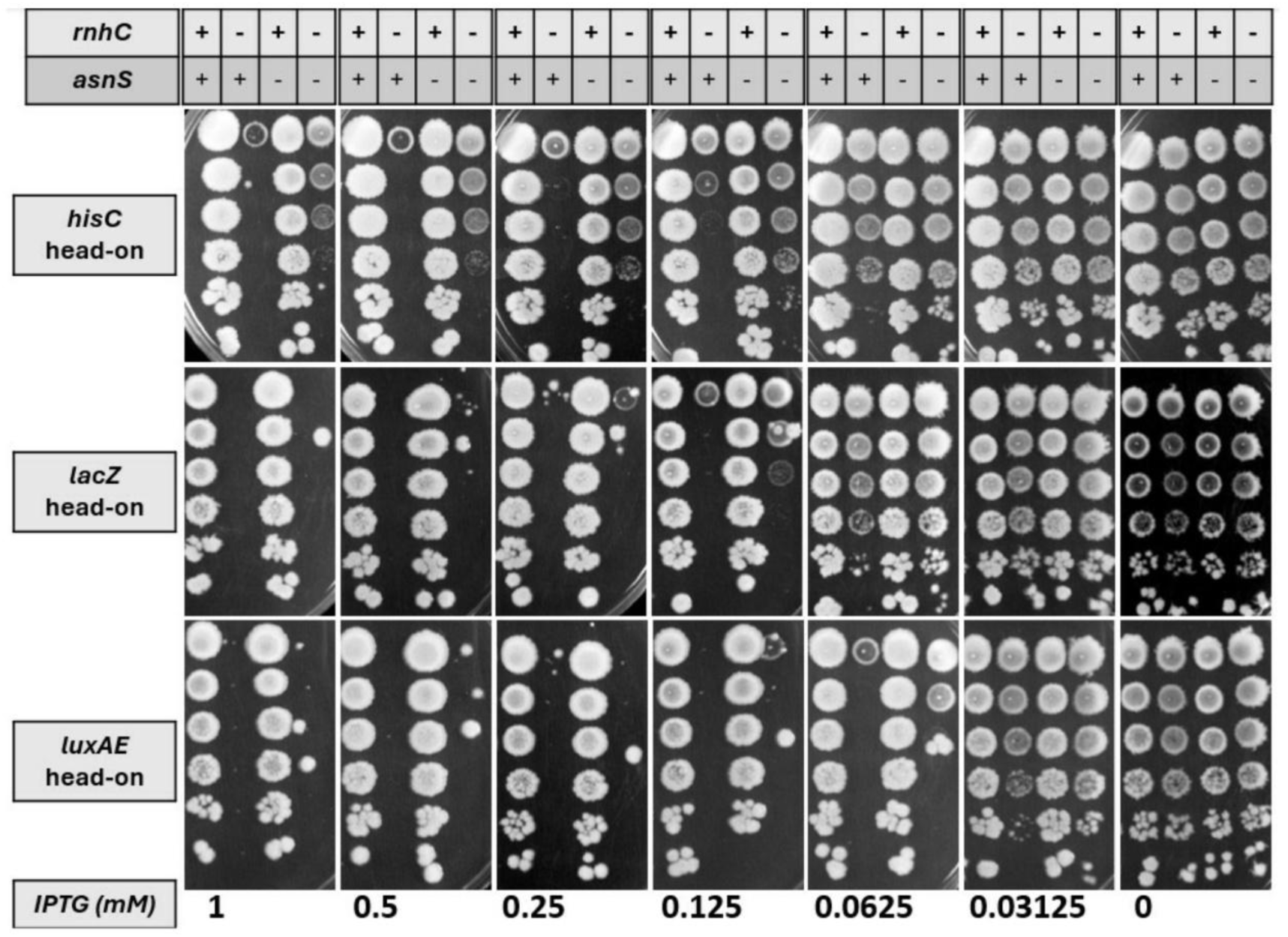
Lethality is reversed in the absence of AsnRS regardless of the gene involved in the engineered conflict. Representative survival plates showing the growth of wild-type, *rnhC*-, *asnS*-, *rnhC*-*asnS*-strains constructed to have the engineered head-on conflict regions involving either of the 3 reporter genes (*hisC*, *lacZ, luxA-E*). Cells were spotted on plates supplemented with various IPTG concentrations [1 - 0 mM]. Plates were incubated at 30°C overnight and removed from the incubator the next morning. Images were taken the following day.

**Supp. Figure 3.**
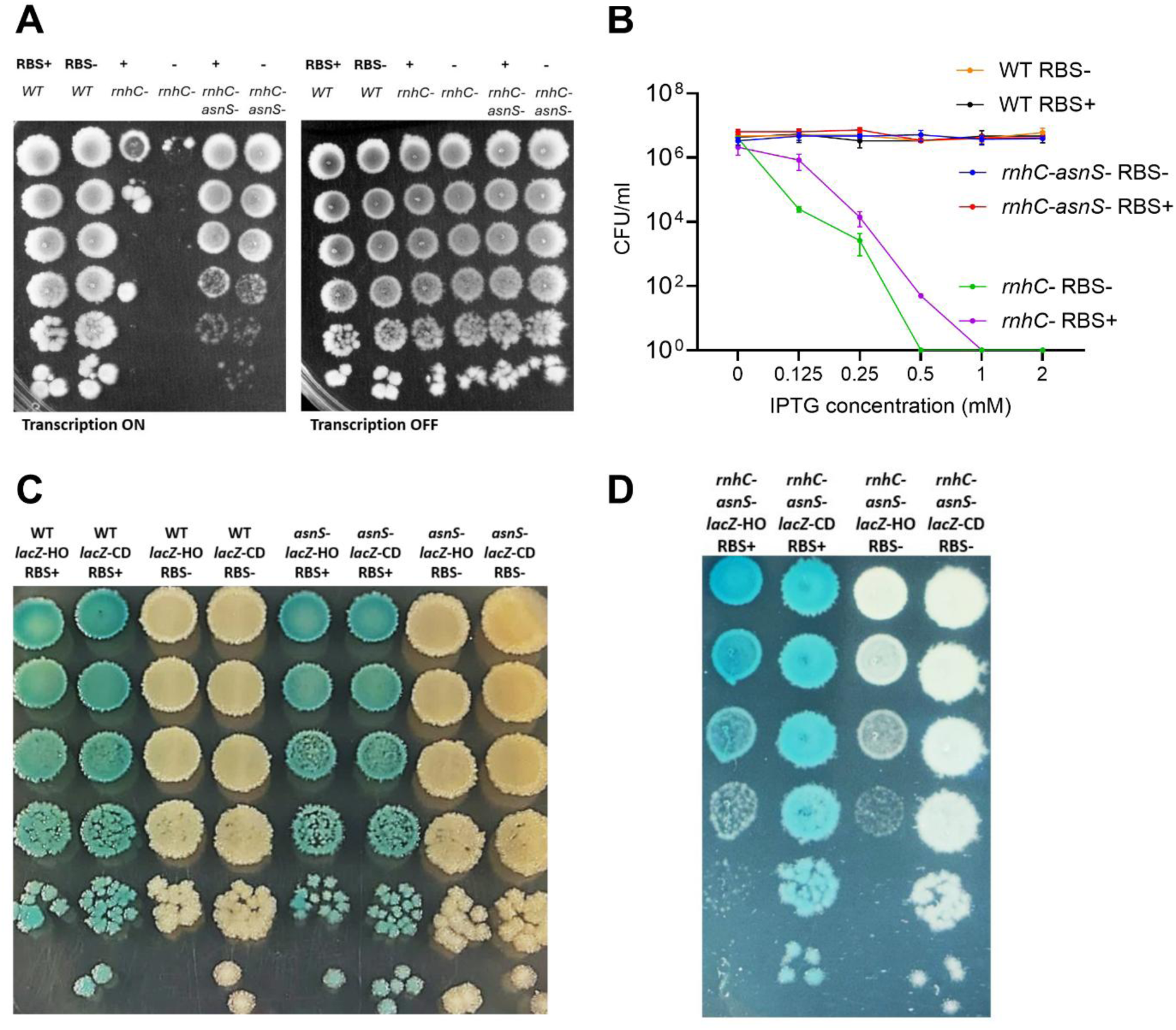
AsnRS role in impairing cell growth does not depend on translation. A) Representative survival plates showing the growth of wild-type, *rnhC*-, *rnhC*-*asnS*-strains that have the reporter gene (*hisC*) with or without its RBS at the engineered head-on conflict region. Cells were spotted on LB plates supplemented with 1mM IPTG (left) and without IPTG (right). After spotting cells, plates were incubated at 30°C overnight and removed from the incubator the next morning. Images were taken the following day. B) Survival assays were performed by inducing the reporter gene expression with IPTG in a dose-dependent manner (x-axis). Survival was measured by calculating CFU/ml for each strain. Data are presented as the mean CFU/ml ± SEM from three independent experiments. C) Survival assays were performed using wild-type or AsnRS-deficient cells constructed to have the engineered conflict regions involving *lacZ* gene, with or without Ribosome Binding Site (RBS), inserted either in head-on or co-directional to replication. The expression of *lacZ* gene was induced with 1mM IPTG and generation of *lacZ* gene product, β-galactosidase, was reported via X-gal (50ug/ml). After spotting cells, plates were incubated at 30°C overnight and removed from the incubator the next morning. Images were taken the following day. D) Survival assays were performed using *rnhC*- and *rnhC- asnS-* cells constructed to have the engineered conflict regions involving *lacZ* gene, with or without Ribosome Binding Site (RBS), inserted either in head-on or co-directional to replication. The expression of *lacZ* gene was induced with 1mM IPTG and generation of *lacZ* gene product, β-galactosidase, was reported via X-gal (50ug/ml). After spotting cells, plates were incubated at 30°C overnight and removed from the incubator the next morning. Images were taken the following day. E) Survival assays were performed using *rnhC*- and *rnhC- asnS-* cells constructed to have the engineered conflict regions involving a non-coding gene, a partial intron of the myc gene from the human genome, inserted either in head-on or co-directional to replication. The expression of the non-coding gene was induced with 0.1mM IPTG. After spotting cells, plates were incubated at 30°C overnight and removed from the incubator the next morning. Images were taken the following day.

**Supp. Figure 4.**
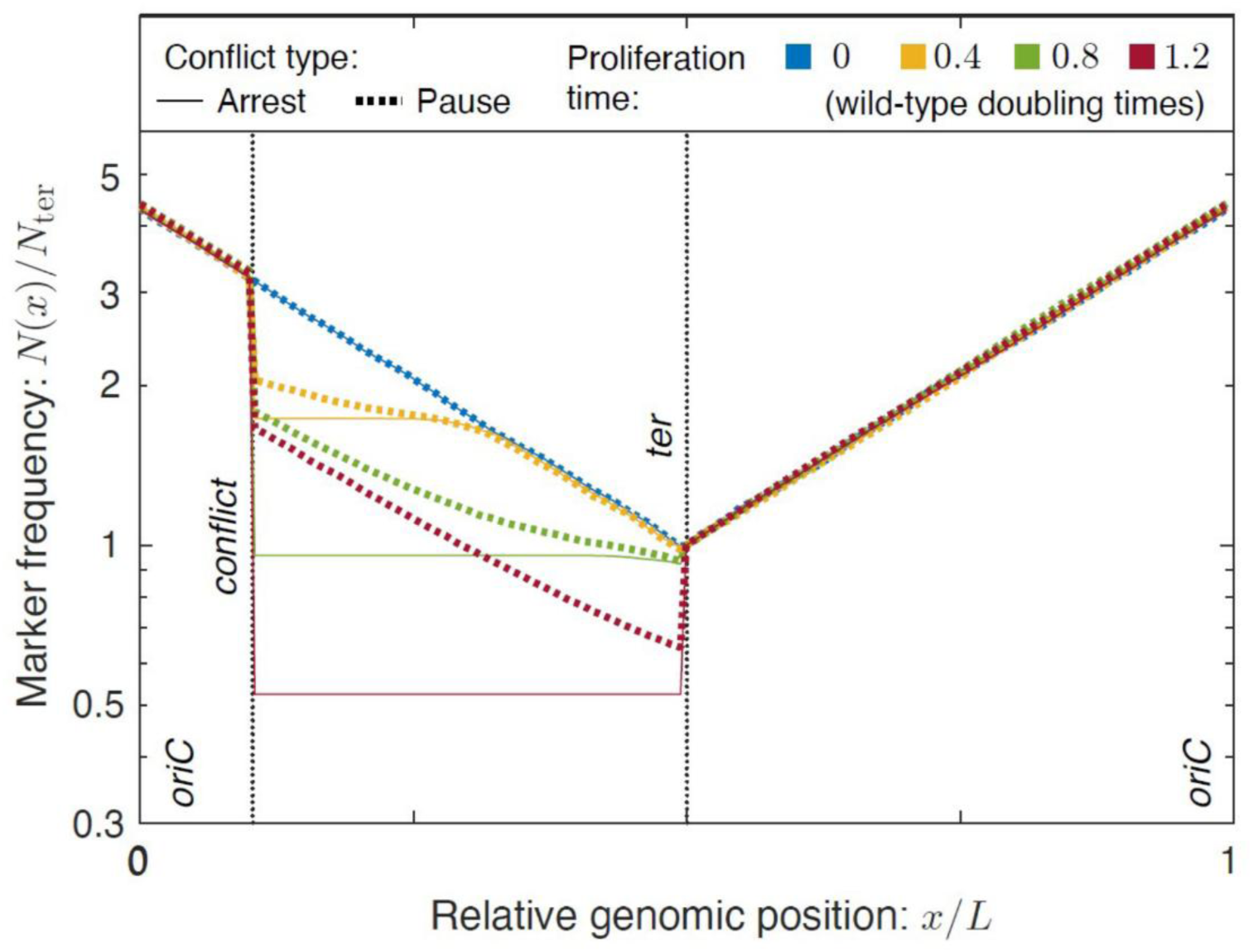
Predicted marker frequencies for induced conflicts. Before conflict induction, the marker frequency is described by the log-phase marker frequency which decays exponential with distance from the origin (blue). Once a conflict is induced, forks are arrested at the conflict location. For arrest-like conflicts, the arrest is perpetual, which leads to the flattening of the marker frequency after the conflict. For pause-like conflicts, the marker frequency has a step-like feature at the conflict, but at long times, the log-slope is constant between the conflict and the terminus as a result of fork motion in this region.

**Supp. Figure 5.**
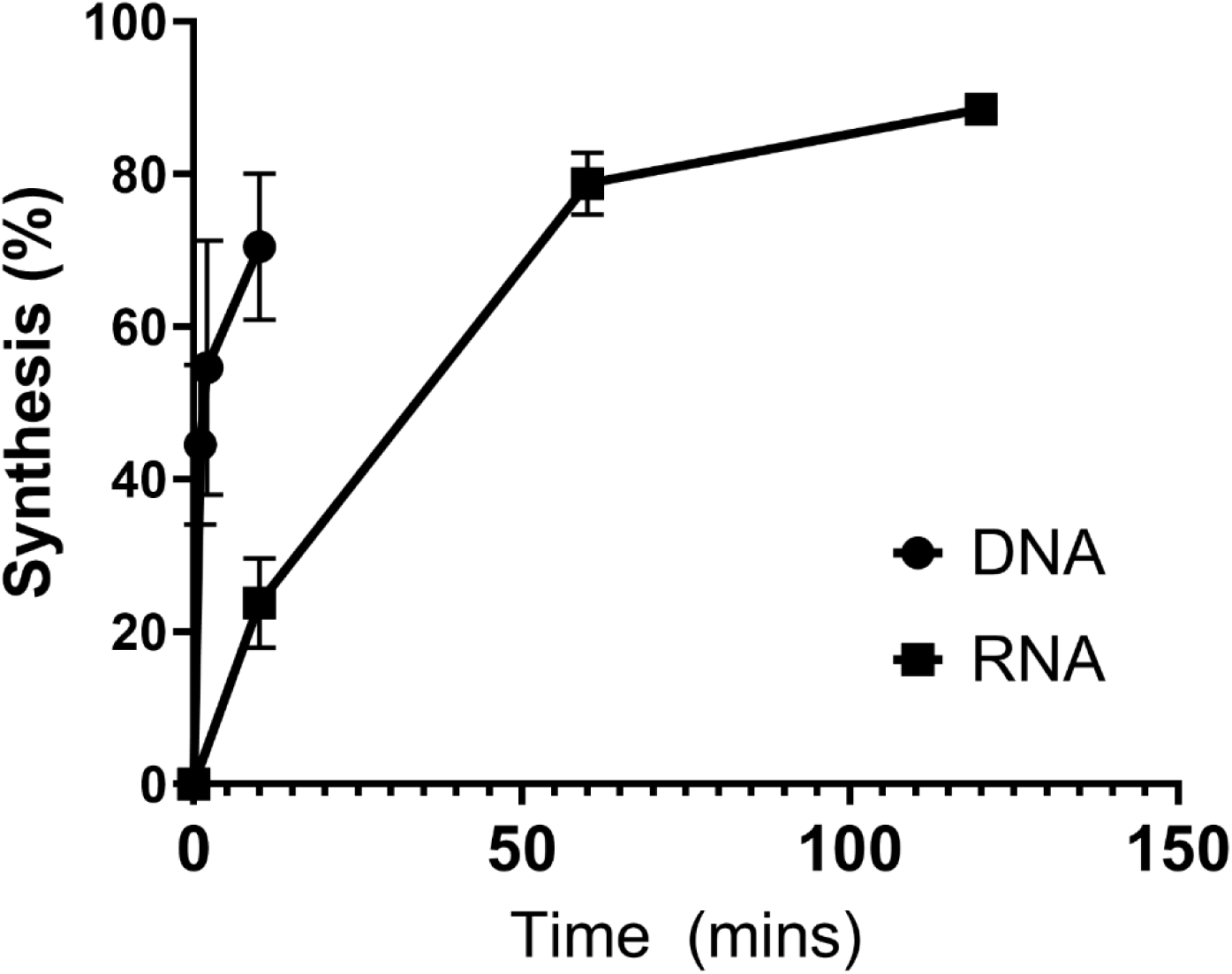
PolA synthesizes DNA from both RNA and DNA primers. *In vitro* primer extension analysis was used to quantify DNA extension by PolA from DNA:DNA and RNA:DNA oligonucleotide templates using dNTPs. Samples were collected at the indicated time points (x-axis), and reactions were stopped using 95% formamide–EDTA. Samples were analyzed on 10% TBE–urea gels. All gels were imaged using a ChemiDoc imaging system (Bio-Rad). The volume of each band was measured by ImageLab software. The percentage synthesis by PolA was calculated by dividing the volume of the top band (extended template) to the total volume of the top and the bottom bands. Data are presented as the mean ± SEM from three independent experiments.

**Supp. Figure 6.**
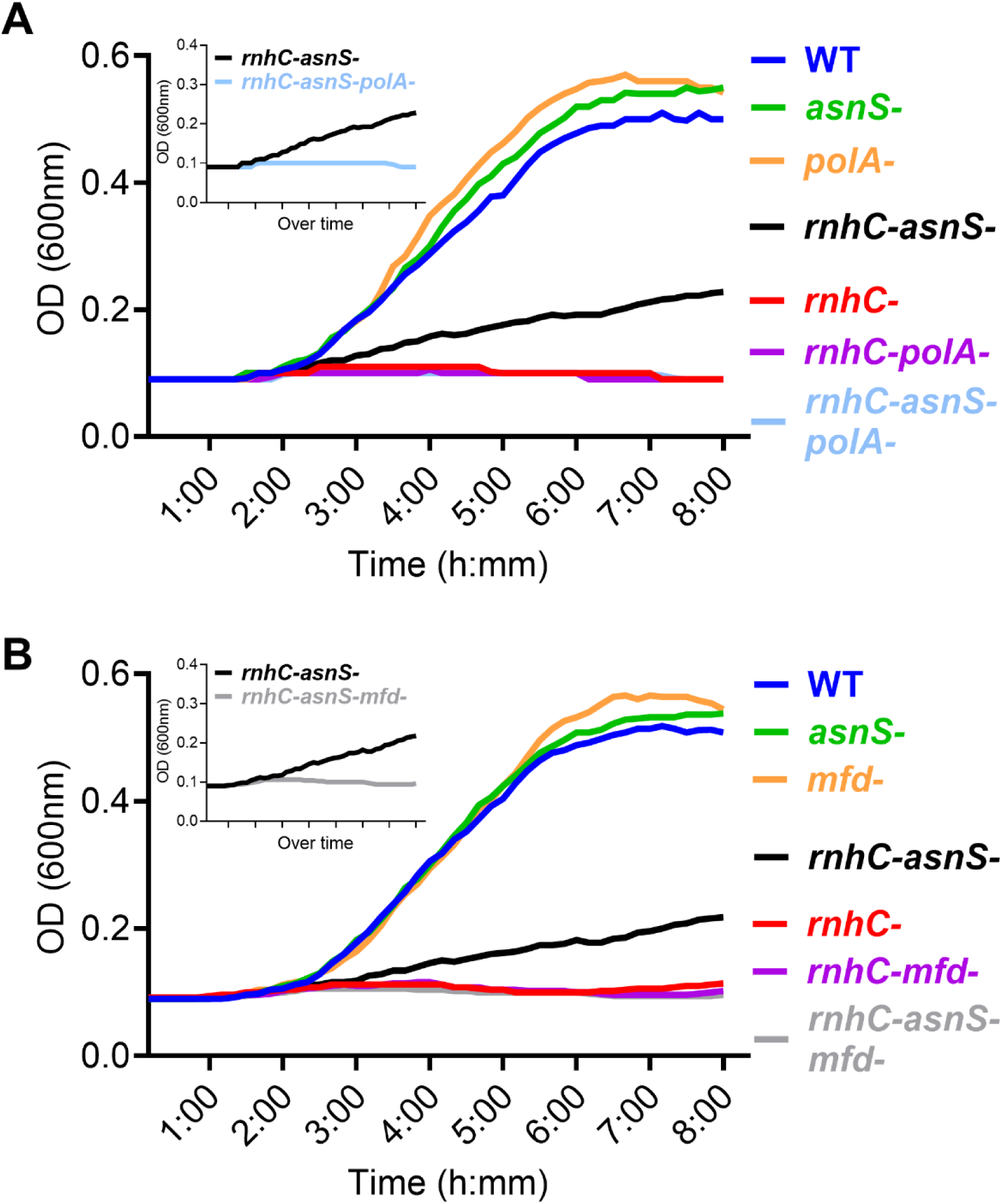
Replication restart from RNA:DNA hybrids at head-on conflict regions requires both Mfd and PolA. A,B) Growth curves of wild-type, *rnhC*-, *asnS*-, *polA*-, *rnhC*-*asnS*-, *rnhC*-*polA*-, *rnhC*-*asnS*-*polA*- and wild-type, *rnhC*-, *asnS*-, *mfd*-, *rnhC*-*asnS*-, *rnhC*-*mfd*-, *rnhC*-*asnS*-*mfd*-cells, respectively, expressing the reporter gene (*hisC*) at the engineered head-on conflict regions. Data were plotted using the mean Optical Density (OD) values over 8 hours. Data are presented as the means of OD values from five independent experiments.

**Supp. Figure 7.**
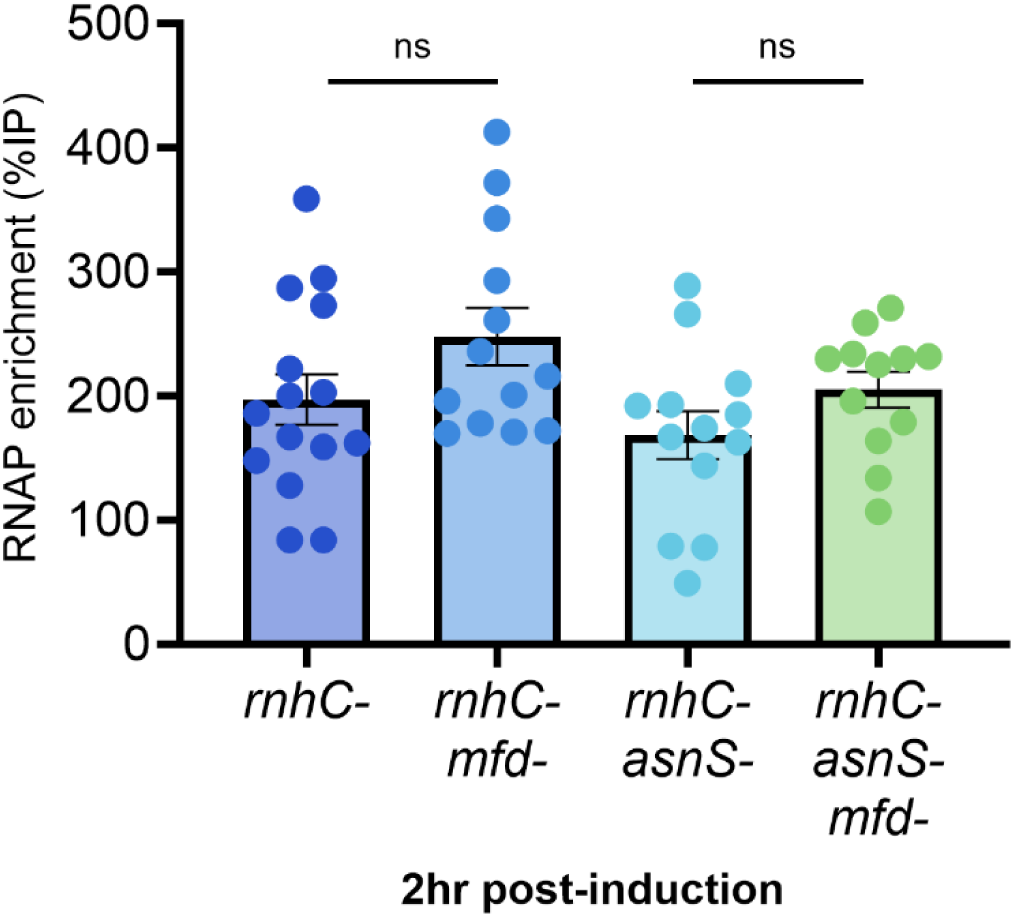
RNAP enrichment at the engineered conflict regions remains similar regardless of the presence/absence of Mfd and AsnRS in cells 2 hours after inducing head-on conflicts. ChIP-qPCR analysis of *rnhC*-*asnS*-, *rnhC*-*asnS*-*mfd*- cells at the early time point, 2 hours after induction of the reporter gene (*hisC*) at the engineered head-on conflict region with IPTG. RNAP was immunoprecipitated using a monoclonal antibody specific to RpoB. Enrichment of RNAP at the engineered head-on conflict regions was quantified by qPCR using %IP method. Data are presented as the mean ± SEM from three independent experiments (n=9). Statistical analysis was performed using one-way ANOVA (ns: not significant).

**Supp. Figure 8.**
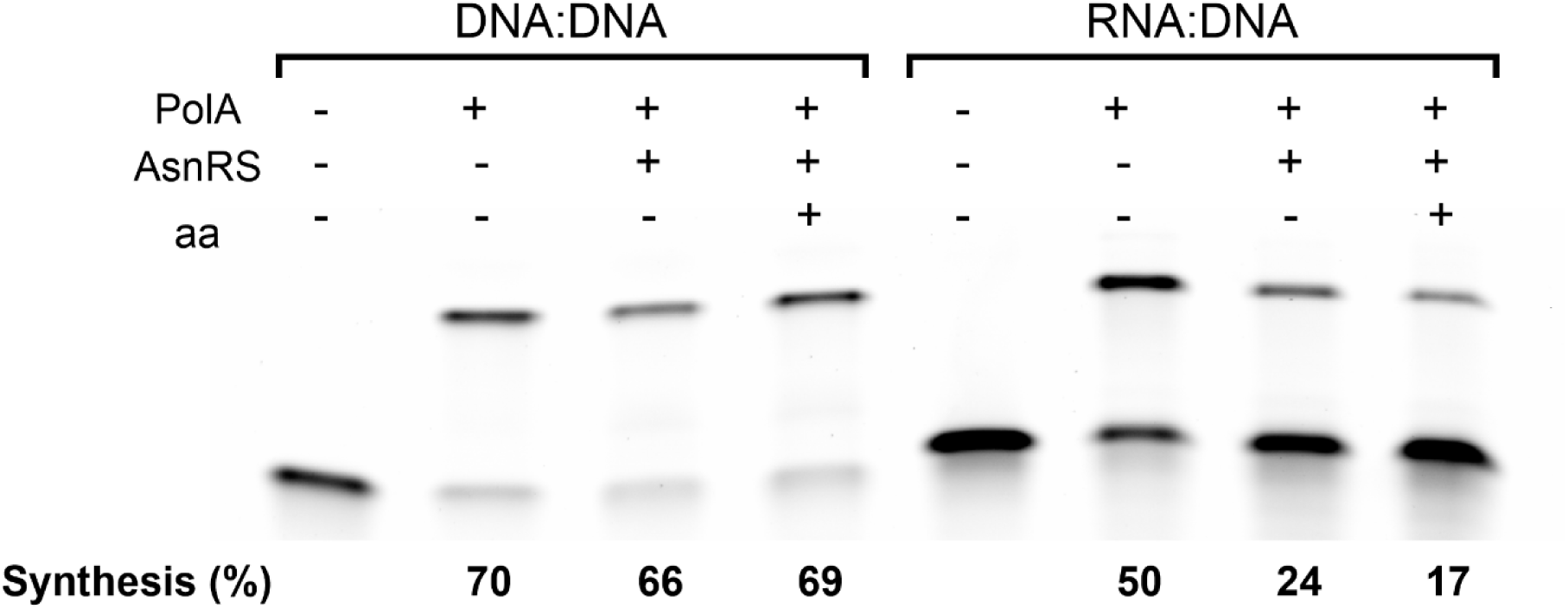
AsnRS does not require asparagine to inhibit PolA function. Representative *in vitro* primer extension analysis quantifying DNA extension by PolA from DNA:DNA and RNA:DNA oligonucleotide templates in the presence and absence of AsnRS and asparagine. Samples were incubated with dNTPs for an hour at 37°C, and reactions were stopped using 95% formamide–EDTA. Samples were analyzed on 10% TBE–urea gels. Volume of each band was quantified using Image lab software and percent synthesis was calculated dividing the top band (extended template) to the whole lane.

**Supp. Figure 9.**
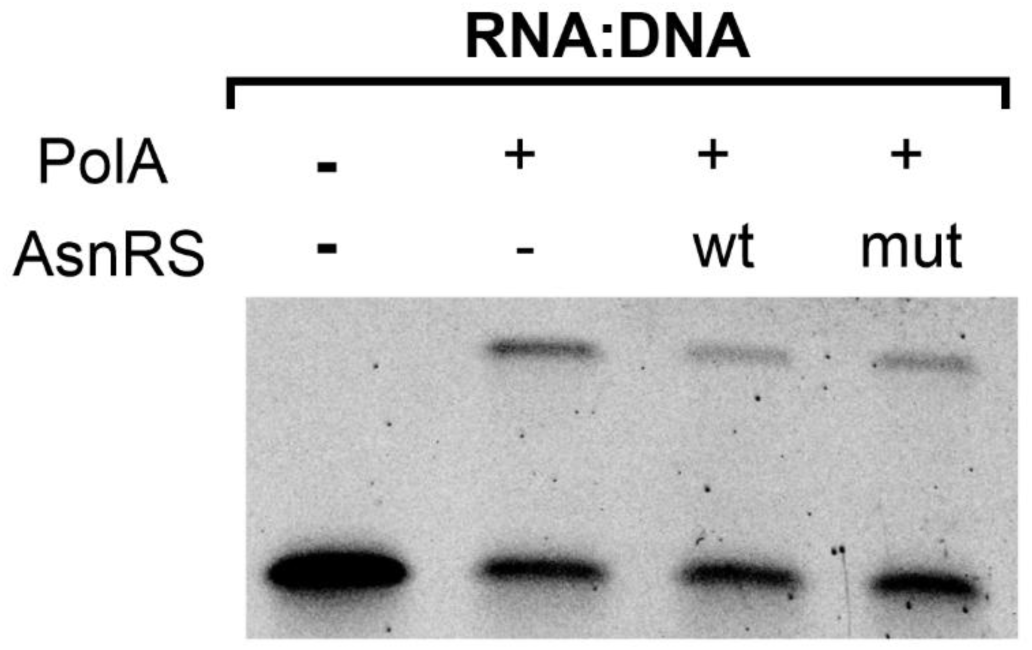
Both AsnRS and AsnRSF219A inhibit DNA extension by PolA from RNA:DNA, but not DNA:DNA template. Representative *in vitro* primer extension analyzing DNA extension by PolA from DNA:DNA and RNA:DNA oligonucleotide templates in the presence and absence of either AsnRS or AsnRSF219A. Samples were incubated with dNTPs for an hour at 37°C, and reactions were stopped using 95% formamide–EDTA. Samples were analyzed on 10% TBE–urea gels.

**Supp. Figure 10.**
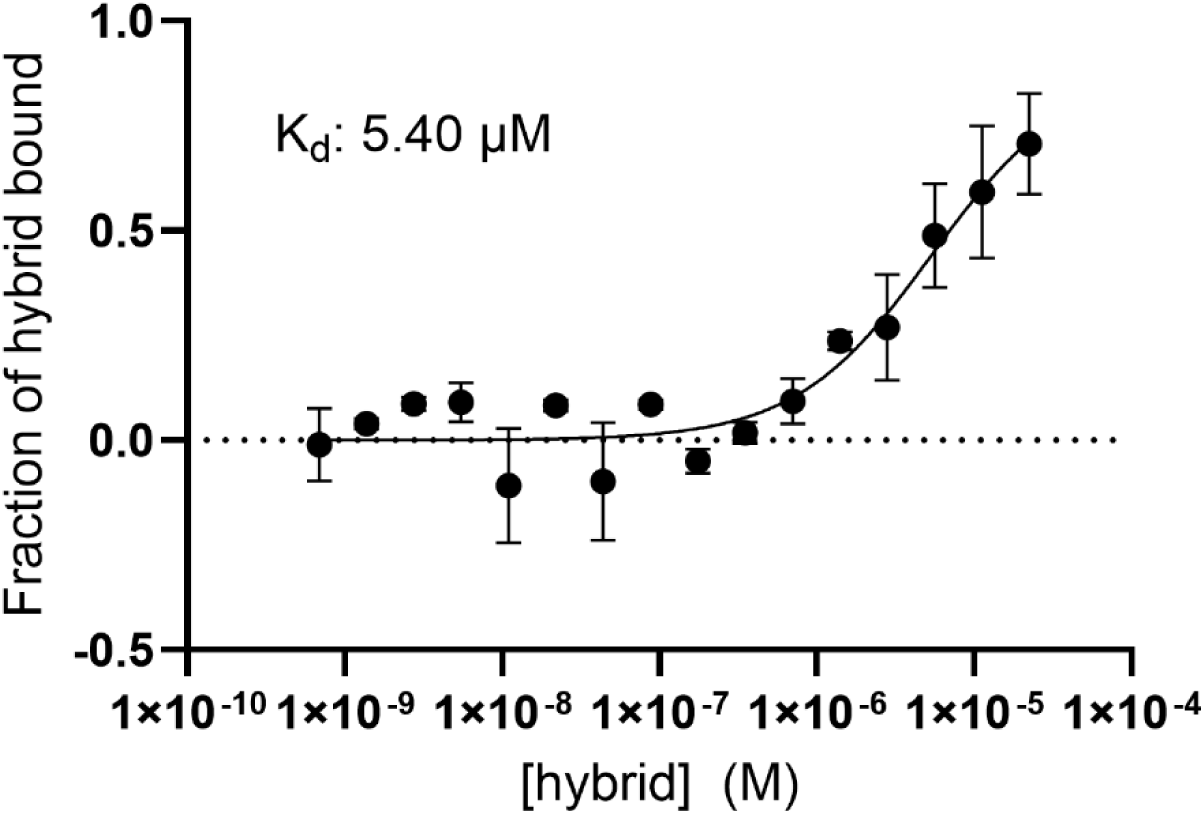
Quantitative analysis of AsnRS binding to an RNA:DNA template by Microscale Thermophoresis. Purified AsnRS was fluorescently labeled and incubated with increasing concentrations of a DNA:RNA hybrid generated by annealed oligonucleotides. Samples were analyzed by MST and binding affinity was determined by fitting the thermophoresis data using GraphPad Prism.

**Supp. Figure 11.**
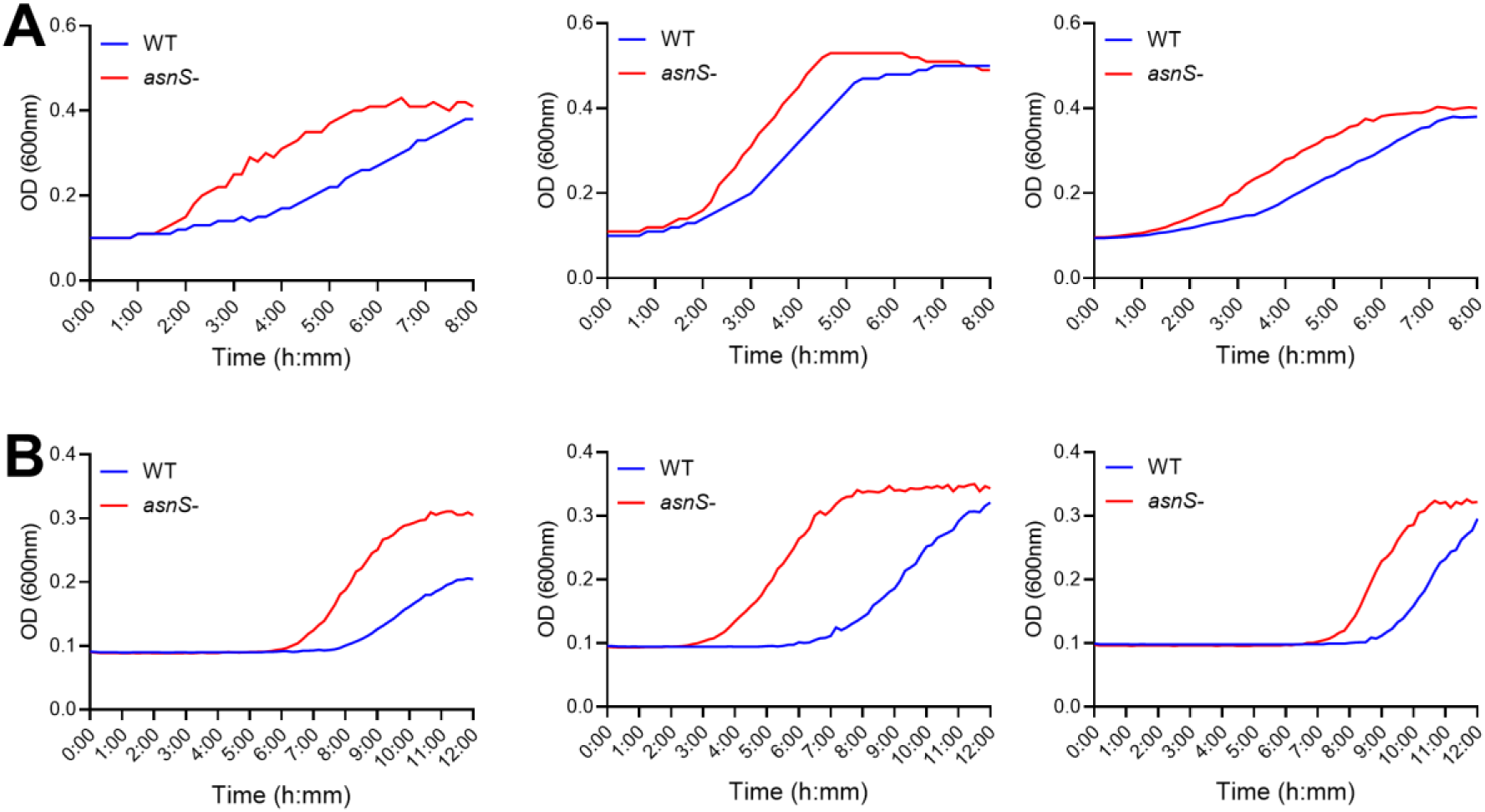
AsnRS regulates replication and growth during exposure to external stresses that induce head-on gene expression. A,B) Growth curves of wild-type and *asnS*- cells in the presence of either 37.5uM paraquat or 2.5ug/ml lysozyme, respectively. Data were plotted using Optical Density (OD) values (y-axis) over 8 and 12 hours (x-axis), respectively. Data are presented as the OD values from three independent experiments for each stressor.

**Supp. Figure 12.**
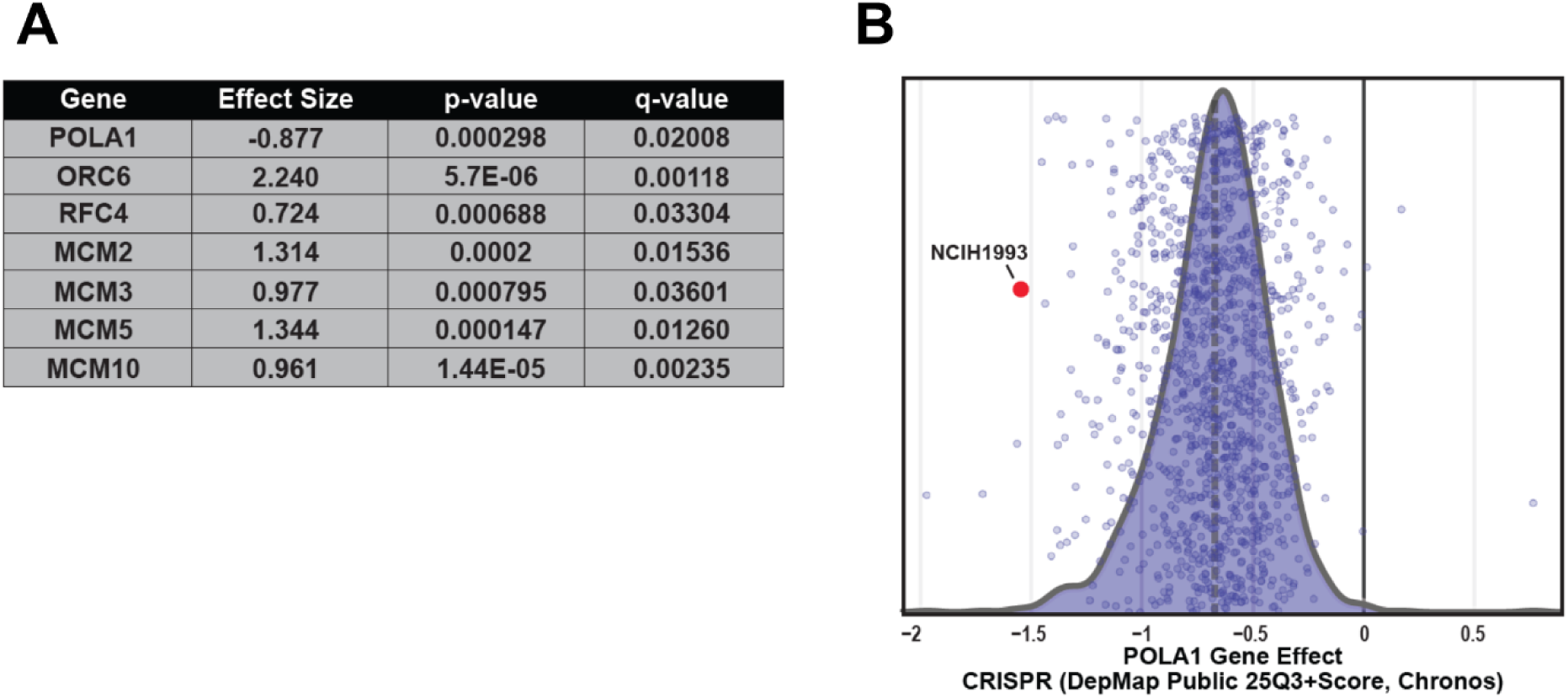
NCI-H1993 cells exhibit enhanced dependency on POLA1. A) Table depicting effect sizes, p-value, and q-value for significant DNA replication-associated genes in NCI-H1993 cells compared to all other cell lines that were identified by Reactome pathway enrichment. B) Distribution of POLA1 gene effect scores from the DepMap Public 25Q3 CRISPR dataset across all profiled cancer cell lines. Individual points represent gene effect values for each cell line. NCI-H1993 highlighted in red.

## Materials and Methods

### Strain Construction

All *B. subtilis* strains were constructed in the HM1 (JH642)^50^ *B. subtilis* background. *B. subtilis* genomic DNA was purified using the GeneJET Genomic DNA Purification Kit (Thermo Scientific) and transformed into HM1 using standard protocols^51^. Strains were streaked on LB agar plates supplemented with selective antibiotics to confirm correct transformation. Additional strain verification was conducted via PCR analysis or Whole Genome Sequencing (Plasmidsaurus). Strains were stored at −80°C in glycerol and used to inoculate experimental pre-cultures. Further details about the strains used in this study are included in Table 1.

**Table 1.** Strains used in this study.

|  |  |  |
| --- | --- | --- |
| 1 | *B. subtilis* | *trpC2 pheA1* |
| 452 | *B. subtilis* | *amyE::Pspank(hy)-hisC (HO) -specR* |
| 1297 | *B. subtilis* | *thrC::Pspank(hy)-lacZ (HO) -mlsR* |
| 1300 | *B. subtilis* | *amyE::Pspank(hy) lacZ (HO) -specR* |
| 1344 | *B. subtilis* | *thrC::Pspank(hy)-lacZ (HO) RBS+* |
| 1366 | *B. subtilis* | *thrC::Pspank(hy)-lacZ (CD) RBS+* |
| 1562 | *B. subtilis* | *thrC::Pspank(hy)-lacZ (HO) RBS-* |
| 1563 | *B. subtilis* | *thrC::Pspank(hy)-lacZ (CD) RBS-* |
| 2043 | *B. subtilis* | *amyE::Pspank(hy)-lacZ (HO) -specR rnhC::mlsR,* |
| 2044 | *B. subtilis* | *amyE::Pspank(hy)-lacZ (CD) rnhC::mlsR,* |
| 2195 | *B. subtilis* | *amyE::Pspank(hy) luxABCDE (HO) -specR* |
| 2655 | *B. subtilis* | *rnhC:: mlsR* |
| 2658 | *B. subtilis* | *amyE::Pspank(hy)-luxABCDE (HO) -specR rnhC:: mlsR* |
| 4257 | *B. subtilis* | *amyE::Pspank(hy)-hisC (HO) -specR rnhC::mlsR* |
| 4680 | *B. subtilis* | *amyE::Pspank(hy)-hisC (HO) RBS+ -specR* |
| 4681 | *B. subtilis* | *amyE::Pspank(hy)-hisC (HO) RBS- -specR* |
| 4682 | *B. subtilis* | *amyE::Pspank(hy)-hisC (HO) RBS+ -specR rnhC::mlsR* |
| 4683 | *B. subtilis* | *amyE::Pspank(hy)-hisC (HO) RBS- -specR rnhC::mlsR* |
| 4734 | *B. subtilis* | *rnhC::markerless amyE::Pspank(hy)-hisC (HO) -specR* |
| 4737 | *B. subtilis* | *asnS::markerless amyE::Pspank(hy)-hisC (HO) -specR* |
| 4740 | *B. subtilis* | *rnhC::markerless asnS::markerless amyE::Pspank(hy)-hisC (HO) -specR* |
| 4748 | *B. subtilis* | *rnhC::markerless asnS::markerless amyE::Pspank(hy)-hisC (HO) -specR*<br>*mfd:: mlsR* |
| 4762 | *B. subtilis* | *rnhC::markerless amyE::Pspank(hy)-hisC (HO) -specR mfd::mlsR* |
| 4804 | *B. subtilis* | *rnhC::markerless asnS::markerless, amyE::Pspank(hy)-hisC (HO) RBS+*<br>*-specR* |
| 4805 | *B. subtilis* | *rnhC::markerless asnS::markerless, amyE::Pspank(hy)-hisC (HO) RBS- -*<br>*specR* |
| 4838 | *E. coli* | pET28a-AsnRS -kanR |
| 4841 | *E. coli* | pET28a-AsnRS-F219A - kanR |
| 4868 | *B. subtilis* | *rnhC*::markerless *asnS*::markerless *amyE*::Pspank(hy)-*hisC* (HO) -*specR*<br>*thrC*::Pspank(hy)-*asnS*(WT) -*mlsR* |
| 4871 | *B. subtilis* | *rnhC*::markerless *asnS*::markerless *amyE*::Pspank(hy)-*hisC* (HO) -*specR*<br>*thrC*::Pspank(hy)-*asnS*(F219A) -*mlsR* |
| 4885 | *B. subtilis* | *rnhC*::markerless *amyE*::Pspank(hy)-*hisC* (HO) -*specR* *polA*::kanR |
| 4887 | *B. subtilis* | *rnhC*::markerless *asnS*::markerless *amyE*::Pspank(hy)-*hisC* (HO) -*specR*<br>*polA*:: kanR |
| 4922 | *B. subtilis* | *amyE*::Pspank(hy)-*hisC* (HO) -*specR* *mfd*::*mlsR* |
| 4923 | *B. subtilis* | *amyE*::Pspank(hy)-*hisC* (HO) -*specR* *polA*::*mlsR* |
| 5062 | *B. subtilis* | *thrC*::Pspank-*lacZ* (HO) RBS+ *asnS*::kanR |
| 5063 | *B. subtilis* | *thrC*::Pspank-*lacZ* (CD) RBS+ *asnS*::kanR |
| 5064 | *B. subtilis* | *thrC*::Pspank-*lacZ* (HO) RBS- *asnS*::kanR |
| 5065 | *B. subtilis* | *thrC*::Pspank-*lacZ* (CD) RBS- *asnS*::kanR |
| 5068 | *B. subtilis* | *rnhC*::markerless *thrC*::Pspank-*lacZ* (HO) RBS+ |
| 5069 | *B. subtilis* | *rnhC*::markerless *thrC*::Pspank-*lacZ* (CD) RBS+ |
| 5070 | *B. subtilis* | *rnhC*::markerless *thrC*::Pspank-*lacZ* (HO) RBS- |
| 5071 | *B. subtilis* | *rnhC*::markerless *thrC*::Pspank-*lacZ* (CD) RBS- |
| 5076 | *B. subtilis* | *rnhC*::markerless *thrC*::Pspank-*lacZ* (HO) RBS+ *asnS*::kanR |
| 5077 | *B. subtilis* | *rnhC*::markerless *thrC*::Pspank-*lacZ* (CD) RBS+ *asnS*::kanR |
| 5078 | *B. subtilis* | *rnhC*::markerless *thrC*::Pspank-*lacZ* (HO) RBS- *asnS*::kanR |
| 5079 | *B. subtilis* | *rnhC*::markerless *thrC*::Pspank-*lacZ* (CD) RBS- *asnS*::kanR |
| 5082 | *B. subtilis* | *rnhC*::markerless *asnS*::markerless *amyE*::Pspank-*hisC* (HO)<br>*thrC*::Pspank-NARS1 |
| 5083 | *B. subtilis* | *rnhC*::markerless *asnS*::markerless *amyE*::Pspank-*hisC* (HO)<br>*thrC*::Pspank-NARS2 |
| 5084 | *B. subtilis* | *asnS*::markerless *amyE*::Pspank-*hisC* (HO) *thrC*::Pspank-NARS1 |
| 5085 | *B. subtilis* | *asnS*::markerless *amyE*::Pspank-*hisC* (HO) *thrC*::Pspank-NARS2 |

### Proteomic Screen

#### DNA:RNA hybrid immunoprecipitation Mass Spectrometry (DRIP-MS)

DRIP-MS samples were collected by optimizing an immunoprecipitation protocol based on modified versions of the DRIP and ChIP assays described below. *B. subtilis* strains were grown on LB agar plates supplemented with selective antibiotics overnight at 37°C. Pre-cultures (3 ml) were grown from a single colony to the mid-exponential phase (OD600 of 0.4-0.7) at 37°C with shaking (260 rpm). To generate a large enough quantity of cells to collect both DNA and proteins from the same population of cells, this culture was first diluted back to OD600 of 0.05 in LB medium (25 ml) and grown to the mid-exponential phase and again diluted back to OD600 of 0.05 in LB medium (100 ml) and grown to the mid-exponential phase at 37°C with shaking (260 rpm). At OD600 of 0.1, the 100 ml cultures were split into two 50 ml cultures. Final concentration of 1mM IPTG was added to one culture to induce transcription at the engineered conflict regions. No IPTG was added to the second culture to create a ‘no transcription’ (no conflict) control sample. Both cultures were crosslinked with 1% formaldehyde for 20 minutes and then quenched with 0.5 M glycine. Crosslinked cells were pelleted by centrifugation (5000 rpm for 5 minutes), washed in cold PBS buffer (20ml), resuspended in 3 ml of Solution A (10 mM Tris-HCl pH 8, 20% w/v sucrose, 50 mM NaCl, 10 mM EDTA, 10 mg/ml lysozyme, 1mM AEBSF), and lysed at 37°C for 30 minutes. 3 ml of 2X IP Buffer (100 mM Tris pH 7, 10 mM EDTA, 20% Trition X-100, 300 mM NaCl, 1 mM AEBSF) was added into lysates and incubated on ice for 30 minutes. Chromosomal DNA was then sheared via sonication (4X 30% amplitude, 10 seconds sonication & 10 seconds rest). Lysates were pelleted by centrifugation (8000 rpm for 15 minutes at 4°C). Lysate supernatant was collected and split into three sub-samples. For INPUT sub-samples, 40 μl was mixed with 40 μl TE Buffer (pH 8) and 9 μl of 10% SDS. For IP sub-samples, lysates were either pre-treated with RNase H enzyme overnight at 37°C (1 ml IP sample) or immediately incubated (5 ml IP sample) overnight at 4°C on a mini-tube rotator (10 rpm) with S9.6 antibody (10 μl/ml IP) (Vanderbilt Antibody and Protein Resource (VAPR) Core). Pre-washed 50% Protein A sepharose beads (GE) (30 μl) were added to IP samples and incubated for 1 hour at room temperature on a mini-tube rotator (10 rpm). Beads were pelleted (2000 rpm for 1 minute) and washed six times for 3 minutes at room temperature in 1X IP Buffer and one time in TE Buffer (pH 8). DNA was eluted off of the beads with Elution Buffer I (100 μl/1ml IP: 50 mM Tris pH 8, 10 mM EDTA, 1% SDS) incubated for 10 minutes at 65°C. Beads were pelleted by centrifugation (5000 rpm for 1 minute) and the supernatant was collected and saved. DNA was eluted off of the beads a second time with Elution Buffer II (150 μl/1ml IP: 10 mM Tris pH 8, 1 mM EDTA, 0.67% SDS). Beads were pelleted again by centrifugation (7000 rpm for 1 minute) and the supernatant was collected and saved with the previous elution. The combined eluates were de-crosslinked at 65°C overnight.

To collect proteins, ∼1000 μl of IP, ∼180 μl of RNase H treated IP, and ∼45 μl of INPUT samples were precipitated one ice by the addition of first 0.11 volumes of cold 100% TCA and then 500 μl of cold 10% TCA. Precipitated proteins were pelleted by centrifugation (20000 rpm for 30 minutes). Protein pellets were washed twice in cold acetone and then air dried before resuspension in 50 μl of a 1:1 H2O 4X SDS Sample Buffer mixture. Protein samples were segregated by size (1 hour at 100 volts) on a 12% Mini Protean TGX SDS-PAGE gel. To visualize proteins, gels were washed three times in water, stained for two hours with Imperial stain, washed three times in water, and then de-stained overnight in water. Gels were then submitted to the Mass Spectrometry Core at Vanderbilt University for LC-MS analysis.

### HPLC-MS

Total protein from each sample was run on an individual lane of an SDS-PAGE gel. The entire lane, except for the two bands corresponding to the light and heavy chain of the antibody, were excised. Each excised lane was saturated with trypsin (10 ng/μl) and allowed to be absorbed at room temperature prior to being digested overnight at 37°C using a standard in-gel digestion protocol. Peptides were extracted from the digested product with two applications of a 60% Acetonitrile / 0.1% Trifluoroacetic acid extraction buffer. Dried pellets were resuspended in 20 μl of 0.2% Formic Acid. Peptide samples (2 μl) were injected into SAM HPLC-MS instrument and run on a 90-minute gradient.

### Analysis and Filtration of DRIP-MS Data

Wild-type and *rnhC*- strains with no other genetic modifications as well as strains containing either *hisC, lacZ, luxA-E* engineered conflict (head-on or co-directional orientation to replication) either with or without *rnhC* gene were used in this experiment. These strains were either treated with or without IPTG which controls transcription.

Raw data received from Vanderbilt MSRC Proteomics Laboratory was visualized using Scaffold 5, proteome software. A list of proteins met the following criteria corresponding to each strain was obtained (Supp. File4_DRIP-MS_1):

Required minimum number of peptides: 2
Target Peptide False Discovery Rate: 0.05
Target Protein False Discovery Rate: 0.05

We then calculated fold change (FC) for each protein by dividing the total spectrum count in IPTG+ to IPTG- pair for each biological replicate. Proteins with FC ≥ 1.5 were selected for each strain. Among those proteins, we selected the ones associated only with head-on oriented engineered conflicts in the absence of *rnhC* gene, subtracting proteins associated with all other strains. After this analysis, we identified a single protein, AsnRS, which was associated with all three conflict regions in the head-on orientation in the absence of RNase HIII, but no other strain.

The follow-up DRIP-MS experiments in cells experiencing head-on conflicts in the absence of RNase HIII with and without AsnRS and/or Mfd were performed following the same protocol. Raw data for those experiments were received from Vanderbilt MSRC Proteomics Laboratory and visualized using Scaffold DDA Proteomics Software. A list of proteins met the following criteria corresponding to each strain was obtained (Supp. File5_DRIP-MS_2):

Required minimum number of peptides: 2
Target Peptide False Discovery Rate: 0.02
Target Protein False Discovery Rate: 0.02

Fold Change calculation was done in the same way as in the initial DRIP-MS screen by applying the same FC ≥ 1.5 threshold to identify ‘enriched’ proteins at the head-on conflict regions only when transcription is induced with IPTG (IPTG+/-).

### Survival Assay

*B. subtilis* strains were grown on LB agar plates supplemented with selective antibiotics at 37°C overnight. Pre-cultures (2 ml) were grown from a single colony to the mid-exponential phase (OD600 of 0.4-0.7) at 37°C with shaking (260 rpm). Pre-cultures were diluted back to OD600 of 0.3 and serially diluted (1 :10) in 1X Spizizen’s Minimal Salts solution. 5μl of each dilution was plated onto LB agar plates supplemented with various concentrations of IPTG (0-1 mM). For x-gal exposure, plates were prepared with both X-gal (80ug/ml) and IPTG (0-1mM). All plates were incubated overnight at 30°C. Plates were imaged with a BioRad Gel Doc XR+ Molecular Imager and colonies were counted. Colony forming units (CFUs) per ml was calculated for each sample at each concentration of IPTG. A minimum of three biological replicates were averaged to calculate the final CFU/ml for each sample-IPTG concentration pair.

### DNA:RNA hybrid immunoprecipitation assays (DRIP)

DRIP assays were performed as previously described^52^ with minor modifications^53^. *B. subtilis* strains were grown on LB agar plates supplemented with selective antibiotics at 37°C overnight. Pre-cultures (3 ml) were grown from a single colony to the mid-exponential phase (OD600 of 0.4-0.7) at 37°C with shaking (260 rpm), diluted back to OD600 of 0.05 in LB medium (25 ml), and grown to the mid-exponential phase again at 30°C while adding 1mM IPTG at an OD600 of 0.1 to induce transcription. Cells reaching mid-exponential phase were then pelleted by centrifugation, washed in cold PBS buffer, and resuspended in TE Buffer (pH 8) supplemented with lysozyme (10 mg/ml). Cell pellets were lysed at 37°C for 30 minutes. Lysis was suspended with the addition of 20% SDS (wt/vol). Lysates were treated with Proteinase K (20 mg/ml) at 37°C overnight. DNA was extracted from lysates with phenol-chloroform-isoamyl alcohol (25:24:1) and precipitated from the aqueous phase with the addition of ethanol (100%) and 3M NaOAc. DNA was spooled, washed twice with ethanol (80%), and air dried at room temperature. DNA was resuspended in TE Buffer (pH 8) and then digested overnight at 37°C with HindIII, EcoRV, EcoRI, and DraI restriction enzymes (NEB). Digested chromosomal DNA was then extracted, with phenol chloroform-isoamyl alcohol (25:24:1), precipitated with ethanol (80%) and 3M NaOAc, resuspended with TE Buffer (pH 8). A DNA subsample (1/10 volume) was collected as INPUT for downstream qPCR analysis. The remaining DNA (IP samples) were either pre-treated with RNase H enzyme overnight at 37°C or immediately incubated overnight at 4°C on a mini-tube rotator (10 rpm) in 10X DRIP Binding Buffer (100 mM NaPO4 pH 7, 1.4 M NaCl, 0.5% Triton X-100) with 20μl of the S9.6 antibody (Vanderbilt Antibody and Protein Resource (VAPR) Core). Pre-washed 50% Protein A sepharose beads (GE) (30μl) were added to IP samples and incubated at 4°C for 2 hours on a mini-tube rotator (10 rpm). Beads were pelleted (2000 rpm for 1 minute) and washed twice in 1X DRIP Binding Buffer for 15 minutes at room temperature on a mini-tube rotator (10 rpm). DNA was eluted off of the beads with 300 μl of DRIP Elution Buffer (10 mM Tris pH 8, 1mM EDTA, 0.67% SDS) and samples were incubated at 55°C for 45 minutes. Beads were pelleted (2000 rpm for 1 minute) and DNA was extracted from the supernatant with phenol-chloroform-isoamyl alcohol (25:24:1) and precipitated with ethanol (80%), 3M NaOAc and glycogen. Pelleted DNA was air dried and then resuspended in 10μl of RNase-free TE Buffer (pH 8). Samples were then analyzed via qPCR and %IP values were calculated for each strain (details below).

### Chromatin immunoprecipitation assays (ChIP)

ChIP assays were performed as previously described^38^ with minor modifications. *B. subtilis* strains were grown on LB agar plates supplemented with selective antibiotics at 37°C overnight. Pre-cultures (3ml) were grown from a single colony to the mid-exponential phase (OD600 of 0.4-0.7) at 37°C with shaking (260 rpm), diluted back to OD600 of 0.05 in LB medium (25ml) with the addition of 1mM IPTG. Cultures were crosslinked with 1% formaldehyde for 20 minutes and then quenched with 0.5 M glycine. Crosslinked cells were pelleted by centrifugation (5000 rpm for 5 minutes at room temperature), washed in ice cold PBS buffer. Pellets were resuspended in 1.5 ml of Solution A (10 mM Tris-HCl pH 8, 20% w/v sucrose, 50 mM NaCl, 10 mM EDTA, 10 mg/ml lysozyme, 1mM AEBSF), and lysed at 37°C for 30 minutes. 3 ml of 2X IP Buffer (100 mM Tris pH 7, 10 mM EDTA, 20% Trition X-100, 300 mM NaCl, 1 mM AEBSF) was added into lysates and incubated on ice for 30 minutes. Chromosomal DNA was then sheared via sonication (4X 30% amplitude, 10 seconds sonication & 10 seconds rest). Lysates were pelleted by centrifugation (8000 rpm for 15 minutes at 4°C). Lysate supernatant was collected and split into two sub-samples. For INPUT sub-samples, 40 μl was mixed with 40 μl TE Buffer (pH 8) and 9 μl of 10% SDS. For IP sub-samples, 1 ml of lysate was incubated with the corresponding antibody on a mini-tube rotator (10 rpm) at 4°C overnight. Pre-washed 50% Protein A sepharose beads (GE) (30 μl) were added to IP samples and incubated for 1 hour at room temperature on a mini-tube rotator (10 rpm). Beads were pelleted (2000 rpm for 1 minute) and washed six times for 3 minutes at room temperature in 1X IP Buffer and one time in TE Buffer (pH 8). DNA was eluted off of the beads with Elution Buffer I (100 μl/1ml IP: 50 mM Tris pH 8, 10 mM EDTA, 1% SDS) incubated for 10 minutes at 65°C. Beads were pelleted by centrifugation (5000 rpm for 1 minute) and the supernatant was collected and saved. DNA was eluted off of the beads a second time with Elution Buffer II (150 μl/1ml IP: 10 mM Tris pH 8, 1 mM EDTA, 0.67% SDS). Beads were pelleted again by centrifugation (7000 rpm for 1 minute) and the supernatant was collected and saved with the previous elution. The combined eluates were de-crosslinked at 65°C overnight.

IP and INPUT samples were diluted in 250 μl of TE Buffer (pH 7.6) and treated with Proteinase K (0.4 mg/ml) at 37°C for 2 hours. DNA was extracted with phenol chloroform-isoamyl alcohol (25:24:1) and chloroform and precipitated with 3M NaOAc, ethanol (100%) and glycogen, and washed with ethanol (75%). Pelleted DNA was air dried and then resuspended in 40 μl of RNase-free TE Buffer (pH 8). Samples were then analyzed via qPCR and %IP values were calculated for each strain (details below).

### Transcript level measurement

*B. subtilis* strains were grown on LB agar plates supplemented with selective antibiotics at 37°C overnight. Pre-cultures (3ml) were grown from a single colony to the mid-exponential phase (OD600 of 0.4-0.7) at 37°C with shaking (260 rpm), diluted back to OD600 of 0.05 in LB medium (25ml) with the addition of 1mM IPTG. Cells were fixed and transcription was halted using 1:1 cold 100% methanol. For total mRNA measurement, total RNA was extracted and purified from 2 ml of cells using the GeneJET RNA Purification Kit (Thermo Scientific). RNA concentration was measured using Qubit™ RNA Broad Range (BR) (Invitrogen) according to the manufacturer’s protocol. 1ug of each RNA sample was treated with DNase I (Thermo Scientific) and converted to cDNA using the iScript cDNA Synthesis Kit (BioRad) including random primers. Samples were then analyzed via qPCR. Transcript levels were calculated by normalizing the relative enrichment of the reporter locus (*hisC*) to a control locus (*katA*).

### Quantitative PCR (qPCR)

Quantitative PCR reactions were performed using iTaq Universal SYBR Green master mix (BioRad) on a CFX96 Touch Real-Time PCR machine (BioRad). IP DNA (1:3 dilution) and INPUT DNA (1:150 dilution) were used as templates for 20μl PCR reactions with 2 uM primers specific to a reporter locus (*lacZ, hisC, luxABCDE*). Two technical replicates were run for each qPCR reaction. The mean Cq value was calculated from the two technical replicates for each biological replicate, provided the standard deviation between replicates was ≤0.5. Mean Cq values were determined separately for IP and INPUT samples, and percent input (%IP) was calculated. A minimum of three biological replicates were analyzed independently and averaged to obtain the final value for each strain.

### Growth curves

*B. subtilis* strains were grown on LB agar plates supplemented with selective antibiotics at 37°C overnight. Pre-cultures (2 ml) were grown from a single colony to the mid-exponential phase (OD600 of 0.4-0.7) at 37°C with shaking (260 rpm). Pre-cultures were diluted back to OD600 of 0.0025 in total volume of 200μl using fresh LB with and without the addition of IPTG (to a 1mM final concentration) in clear 96-well plates. For stress exposure, instead of IPTG, either paraquat (37.5mM) or lysozyme (2.5ug/ml) was added into the total volume of 200μl. Plates were then incubated in an Epoch2 plate reader set to 37°C with shaking (260rpm) for 8 to 12 hours, taking OD measurements every 10 minutes. Each biological replicate had 2 technical replicates which then were averaged followed by the mean and SEM calculations of desired number of biological replicates per experiment. The data was plotted in GraphPad Prism10.

### Marker frequency analysis

*B. subtilis* strains were grown on LB agar plates supplemented with selective antibiotics at 37°C overnight. Pre-cultures were grown from a single colony to the mid-exponential phase (OD600 of 0.4-0.7) at 37°C with shaking (260 rpm), diluted back to OD600 of 0.05 in LB medium with the addition of 1mM IPTG. Cells were harvested in 1, 2, 3 or 7 hours. DNA was purified using the GeneJET Genomic DNA Purification Kit (Thermo Scientific) and sent to VUMC Technologies for Advanced Genomics Facility for DNA library preparation and PE150 sequencing on the NovaSeq 6000 targeting an average of 8M reads. The reads filtered with fastp^54^ and mapped to the genome of the *B. subtilis* strains tested using Bowtie 2^55^.

The coordinate of first nucleotide of each read was used to plot the normalized read counts on GraphPad Prism10, and a function, smooth curve, was utilized to average 5 neighbors to each side and re-plot to smoothen the curves.

### AsnRS purification

Coding regions of *asnS_Bsub_* and *asnSF219A_Bsub_*, excluding the start codon, was amplified using Q5 polymerase (NEB) and inserted into the pET28a vector (Thermo) between BamHI and XhoI sites to create a construct with an N-terminal 6×His tag. This plasmid was then transformed into Rosetta (DE3) E. coli cells. A pre-culture supplemented with kanamycin was grown from a single colony at 37°C overnight with shaking (260 rpm). The culture was then diluted (1:100) in 250ml of LB with kanamycin and grown at 37°C for 3 hours until it reached an OD600 of 0.2. To induce protein expression, 1 mM IPTG was added, and cells were incubated at 37°C for 4 hours. The purification process was performed similarly to previously described methods^56,57^. Cells were pelleted (at 4000xg for 5 mins) resuspended in in wash buffer (25 mM HEPES, pH 7.2, 250 mM NaCl, 10 mM imidazole, and 5 mM 2-mercaptoethanol) with 0.1% Triton X-100 and 0.5mM AEBSF. The suspension was then sonicated for 5 min total, 2 sec on and 2 s off alternating at 40% amplitude. The homogenate was centrifuged at 21000xg for 45 min at 4°C. The supernatant was run through, twice, HisPur™ Ni-NTA resin (Thermo), that was equilibrated with wash buffer. The column was washed, twice, with the wash buffer and AsnRS was eluted with the wash buffer supplemented with 250mM imidazole. Eluted AsnRS was dialyzed in storage buffer (25 mM HEPES–KOH, pH 7.2, 30 mM KCl, 0.2 mM EDTA, 10% glycerol, and 5 mM 2-mercaptoethanol) at 4°C overnight, concentrated using Amicon Ultra centrifugal filter units, and stored at −80°C. Aliquots collected during protein preparation steps were run on polyacrylamide gel and Coomassie-stained to confirm the purity. Purified and concentrated protein was also detected by Western Blot.

### Primer Extension Assay

PolA activity was assessed^47^ in a reaction buffer (40 mM Tris (pH 8), 10 mM MgCl₂, 60 mM KCl, 2.5% glycerol) with 1 mM dNTPs, 1.5 nM of the designated DNA:DNA or RNA:DNA substrate labeled with Cy5 (IDT), and 100 nM PolA. Reactions (10μl) were incubated at 37°C and samples were collected at 10 minutes, 30 minutes, 1 hour and 2 hours. Reactions were terminated by adding 10μl of 95% formamide containing 10 mM EDTA. DNA was denatured by heating at 85°C for 15 minutes and run on a 12% urea-denaturing polyacrylamide gel (Bio-Rad) at 150 V for 30 minutes in warm 1X TBE Buffer.

AsnRS impact on PolA activity was assessed in the same way, but with or without 100 nM AsnRS (either wild-type or F219A mutant) and 2mM asparagine. Reactions (10μl) were incubated at 37°C and samples were collected 1 hour and terminated by adding 10μl of 95% formamide containing 10 mM EDTA. DNA was denatured by heating at 90°C for 5 minutes and run on a 10% urea-denaturing polyacrylamide gel (Bio-Rad) at 150 V for 25 minutes.

All gels were imaged using a ChemiDoc imaging system (Bio-Rad). The volume of each band was measured by ImageLab software. The percentage synthesis by PolA was calculated by dividing the volume of the top band (extended template) to the total volume of the top and the bottom bands (the entire amount of template added into the reaction).

Templates used in EMSA and Primer extension:

RNA:DNA Duplex sequence:

5’-/5Cy5/rArUrUrCrUrGrGrUrGrGrArArArUrGrGrCrGrCrGrCrUrGrCrUrA -3’
5’-GTGGAACGCTATATGTGCCATTAGCAGCGCGCCATTTCCACCAGAAT -3’

DNA:DNA Duplex sequence:

5’-/5Cy5/ATTCTGGTGGAAATGGCGCGCTGCTA -3’
5’-GTGGAACGCTATATGTGCCATTAGCAGCGCGCCATTTCCACCAGAAT -3’

### Electromobility Shift Assay (EMSA)

AsnRS binding to either DNA:DNA or RNA:DNA templates, labeled with Cy5 (IDT), was assessed in a reaction buffer (50 mM Tris (pH 7.5), 100 mM KCl, 5% glycerol, 1mM EDTA, 10mM DTT), 1.5 nM of the designated substrate, and 100nM of either AsnRS^WT^ or AsnRS^F219A^ with and without 2mM asparagine and 2mM ATP. Reactions (10μl) were incubated at 37°C for 30 minutes. Samples were run on 5% polyacrylamide gel at 4°C, 100 V for 60 minutes.

### Microscale Thermophoresis (MST)

Microscale thermophoresis was performed on a Monolith NT.115 (NanoTemper) as in Carvajal-Garcia et. al. (2024) with some modifications. Purified, his-tagged AsnRS was labeled using the His-Tag Labeling Kit (NanoTemper) according to manufacturer’s protocol. 16 serial 1:2 dilutions of a DNA:RNA hybrid made by annealing oligos, listed below, in a thermocycler were incubated with labeled 20 nM AsnRS on ice for 10 minutes. MST experiments were performed under default settings and GraphPad Prism 10 was used to determine the K_d_.

Templates used in MST:

Bottom RNA strand:
5’-rGrUrGrGrArArCrGrCrUrArUrArUrGrUrGrCrCrArUrUrArGrCrArGrCrGrCrGrCrCrA rUrUrUrCrCrArCrCrArGrArArU
Top DNA strand:
5’-GCGCGCTGCTAATGGCA

### Pyrophosphate Assay

Aminoacylation activity of AsnRS^WT^ and AsnRS^F219A^ was assessed *in vitro* by the addition of 2 mM L-Asn, 2mM ATP, 2uM purified AsnRS, 10ug/ml yeast tRNA (Invitrogen), 1 mM DTT, 1 U/ml inorganic pyrophosphatase in the aminoacylation buffer (30 mM HEPES pH 7.2 containing 140 mM NaCl, 30 mM KCl, and 40 mM MgCl2) at 37°C for 30 min^58,59^. The total reaction volume was 10μl for each sample. Then the manufacturer’s protocol was followed (Sigma, MAK168). The fluorescence measurements were done using BioTek NEO2 with Alpha Screening (Agilent).

### DepMap analysis of NARS1

CRISPR dependency data were obtained from the DepMap Public 25Q3 CRISPR dataset^60,61^. All analyses were performed using DepMap-provided tools unless otherwise specified. To assess global dependency on NARS1 across cell lines, gene effect scores were queried directly in the DepMap portal. The distribution of NARS1 gene effect scores across all profiled cell lines was visualized using DepMap’s built-in plotting tools, and the resulting figure was exported as a scalable vector graphics (SVG) file. To identify genes with differential dependency patterns associated with resistance to NARS1 loss, a two-class comparison was performed in DepMap using the NCI-H1993 cell line as one class and all remaining cell lines as the comparison group. For each gene an effect size was calculated as the difference in mean gene effect between the two classes. Statistical significance was determined as part of DepMap’s built-in analysis pipeline, and false discovery rate (FDR)–adjusted q-values were reported. The top 1,000 genes ranked by statistical significance were visualized in a volcano plot. The volcano plot was exported directly from DepMap as an SVG file for downstream figure assembly. Genes exhibiting significant differential dependency (q < 0.05) in the two-class comparison were subjected to pathway enrichment analysis using Reactome. Enrichment analysis was performed in R, and pathways were ranked based on adjusted p-values and gene ratios. To visualize functional relationships among replication-associated genes identified in the pathway enrichment analysis, a protein-protein interaction network was generated in STRING using default settings^62^.

### Cloning of NARS1 and NARS2

The human NARS1 and NARS2 cDNA sequences (Supp. File3_asnS_cDNA) were obtained from the UCSC Genome Browser. NARS1 and NARS2 were codon-optimized for expression in B. subtilis using the Codon Optimization Tool from Integrated DNA Technologies (IDT). 25 bp overlaps were added to both DNA ends for cloning and the cDNAs were purchased as gene fragments from IDT. NARS1 and NARS2 gene fragments were cloned into HpaI-linearized (NEB # R0105L) pFastBac Dual (Gibco #10712024) using NEBuilder HiFi DNA Assembly Master Mix (NEB #E2621L), transformed into 10-beta competent E. coli (NEB #C3019H). Plasmids were confirmed by whole plasmid sequencing (Azenta Life Sciences).

### Method for simulating bacterial marker frequency

See ‘SuppFile2_Sim_Rep’ and ‘SM1’.

## Acknowledgments

We would like to thank W. Hayes McDonald from Vanderbilt MSRC Proteomics Laboratory and former Merrikh lab member Dan Pers for technical help and support in DRIP-MS experiments, as well as Vanderbilt Technologies for Advanced Genomics (VANTAGE) Core for conducting sequencing experiments. H.M. And O.S. were supported by Vanderbilt University School of Medicine Funds. J.C.G. was supported by HHWF fellowship AND K99ES037493. J.H. was supported by Wayne State University Startup Funds. P.A.W. was supported by NIH grant R01-GM128191 and NSF grant GR046955.

